# Evolutionary transition to XY sex chromosomes associated with Y-linked duplication of a male hormone gene in a terrestrial isopod

**DOI:** 10.1101/2020.05.18.102079

**Authors:** Aubrie Russell, Sevarin Borrelli, Rose Fontana, Joseph Laricchiuta, Jane Pascar, Thomas Becking, Isabelle Giraud, Richard Cordaux, Christopher H. Chandler

**Author notes:** Corresponding author: Christopher H. Chandler Department of Biological Sciences State University of New York at Oswego.

## Abstract

Sex chromosomes are highly variable in some taxonomic groups, but the evolutionary mechanisms underlying this diversity are not well understood. In terrestrial isopod crustaceans, evolutionary turnovers in sex chromosomes are frequent, possibly caused by *Wolbachia*, a vertically-transmitted endosymbiont causing male-to-female sex reversal. Here, we use surgical manipulations and genetic crosses, plus genome sequencing, to examine sex chromosomes in the terrestrial isopod *Trachelipus rathkei*. Although an earlier cytogenetics study suggested a ZZ/ZW sex chromosome system in this species, we surprisingly find multiple lines of evidence that in our study population, sex is determined by an XX/XY system. Consistent with a recent evolutionary origin for this XX/XY system, the putative male-specific region of the genome is small. The genome shows evidence of Y-linked duplications of the gene encoding the androgenic gland hormone, a major component of male sexual differentiation in isopods. Our analyses also uncover sequences horizontally acquired from past *Wolbachia* infections, consistent with the hypothesis that *Wolbachia* may have interfered with the evolution of sex determination in *T. rathkei*. Overall, these results provide evidence for the co-occurrence of multiple sex chromosome systems within *T. rathkei*, further highlighting the relevance of terrestrial isopods as models for the study of sex chromosome evolution.

## Introduction

Although sexual reproduction is shared by most eukaryotes, a variety of different cues can trigger individuals to follow a male, female, or hermaphroditic developmental plan (Conover and Kynard, 1981; Tingley and Anderson, 1986; Janzen and Phillips, 2006; Ospina-Álvarez and Piferrer, 2008; Verhulst *et al*., 2010). In many eukaryotes, sex is primarily determined genotypically, often involving sex chromosomes, although other mechanisms, such as polygenic systems and haplodiploidy, are also known (Vandeputte *et al*., 2007; Heimpel and de Boer, 2008). Sex chromosomes in animals are usually grouped into two main classes: XY systems, in which males are heterogametic (XY) and females are homogametic (XX); and ZW systems, in which females are heterogametic (ZW) and males are homogametic (ZZ). However, non-genetic cues can also play an important role in some species. For instance, environmental factors, such as temperature or population density, influence or determine phenotypic sex in reptiles, fishes, and invertebrates (Conover and Kynard, 1981; Tingley and Anderson, 1986; Janzen and Phillips, 2006). In some cases, cytoplasmic factors, including sex-reversing endosymbionts, such as *Wolbachia*, microsporidia, and paramyxids can serve as a sex-determining signal (Terry *et al*., 1998; Bouchon *et al*., 1998; Kageyama *et al*., 2002; Negri I *et al*., 2006; Pickup and Ironside, 2018).

Evolutionary theory holds that the formation of sex chromosomes begins when an autosome acquires a sex-determining locus (Rice, 1996). Subsequently, recombination around the sex-determining locus is selected against because of sexually antagonistic selection (Bergero and Charlesworth, 2009). For instance, selection should favor mutations that are beneficial in males but deleterious in females when those alleles are linked to a dominant male-determining allele; recombination, on the other hand, would break up this linkage and result in females that carry these male-beneficial alleles. The non-recombining region is then expected to spread in the presence of continued sexually antagonistic selection, and may eventually span the whole sex chromosome, except for the usual presence of a small recombining pseudo-autosomal region (Charlesworth *et al*., 2005). Once recombination has ceased, the non-recombining sex chromosome, such as the Y chromosome in mammals or the W chromosome in birds, is expected to degenerate. Non-recombining genes frequently undergo pseudogenization, acquiring nonsense mutations or transposable element insertions (Charlesworth and Charlesworth, 2000). At the same time, gene trafficking can occur when selection promotes the translocation of formerly autosomal genes to the sex chromosomes (Emerson *et al*., 2004).

Different species appear to be at different stages of sex chromosome evolution. For instance, the sex chromosomes of therian mammals are highly conserved, having originated ∼160 million years ago (Potrzebowski *et al*., 2008; Veyrunes *et al*., 2008). The highly degenerated, heteromorphic Y chromosome represents an advanced stage of sex chromosome evolution. In other taxonomic groups, on the other hand, sex chromosomes appear to undergo more frequent evolutionary turnovers (Ross *et al*., 2009; Cioffi *et al*., 2013; Vicoso and Bachtrog, 2015; Myosho *et al*., 2015; Pennell *et al*., 2018; Jeffries *et al*., 2018). Such young sex chromosomes may have little or no recombination suppression, differentiation in gene content, or sex chromosome dosage compensation, and may not be detectable by traditional cytogenetic methods because they are visually indistinguishable (homomorphic) (Gamble *et al*., 2014; Vicoso and Bachtrog, 2015). Sex chromosomes may even be polymorphic within a species, with different sex-determining loci segregating within or among populations (Orzack *et al*., 1980; Traut, 1994; Ogata *et al*., 2008; Meisel *et al*., 2016).

Unfortunately, we still have a limited understanding of why evolutionary turnovers of sex chromosomes are rare in some groups but frequent in others. A variety of models have been proposed to explain why these turnovers occur, including sexual antagonism, where a novel sex-determining allele spreads because of its association with another allele with sex-specific effects (van Doorn and Kirkpatrick, 2007); the accumulation of deleterious mutations on the nonrecombining sex chromosome (Blaser *et al*., 2013); and the ‘hot potato’ model, which suggests that the accumulation of both sexually antagonistic and deleterious mutations can lead to repeated sex chromosome turnovers (Blaser *et al*., 2014). In some organisms, interactions with vertically transmitted reproductive endosymbionts are also thought to influence the evolution of their hosts’ sex determination mechanisms (Rigaud *et al*., 1997; Cordaux *et al*., 2011). However, many of these models have been difficult to test in nature. This problem is exacerbated by the fact that, while sex chromosomes have been extensively studied in model organisms like *Drosophila*, studies are more sparse in non-model organisms.

One group that has received relatively little attention is crustaceans. Different crustacean species show a variety of distinct sex determining mechanisms, yet there are very few crustacean species in which candidate master sex-determining genes have been identified (Chandler *et al*., 2017, 2018). Within crustaceans, perhaps one of the best-studied groups in terms of sex determination is the terrestrial isopods (Oniscidea). Terrestrial isopod species have a mix of XY and ZW systems, along with reports of a few parthenogenic species and populations (Fussey, 1984; Johnson, 1986; Rigaud *et al*., 1997). The bacterial endosymbiont *Wolbachia* also influences sex determination by causing male-to-female sex reversal in some isopod hosts (Bouchon *et al*., 1998; Cordaux *et al*., 2004). In fact, interactions with *Wolbachia* are thought to drive rapid evolutionary turnover of the sex chromosomes in terrestrial isopods.

For instance, in the common pillbug *Armadillidium vulgare*, a copy of the *Wolbachia* genome horizontally integrated into the host genome (known as the *f* element) led to the origin of a new W chromosome (Leclercq *et al*., 2016). A recent phylogenetic analysis also identified several transitions in heterogametic systems along the isopod phylogeny, including closely related species pairs with different sex chromosome systems (Becking *et al*., 2017). Moreover, only a few species of terrestrial isopods are known to have heteromorphic sex chromosomes, in which the X and Y, or Z and W, chromosomes are distinguishable in cytogenetics experiments (Rigaud *et al*., 1997), and WW or YY individuals are often viable and fertile (Juchault and Rigaud, 1995; Becking *et al*., 2019), suggesting that the W and Y chromosomes have not lost any essential genes in these species.

In this study, we examined sex determination in the widespread species *Trachelipus rathkei.* This species was previously established by cytogenetic methods to have heteromorphic, albeit slightly, Z and W sex chromosomes (Mittal and Pahwa, 1980), and is nested within a clade that appears ancestrally to possess a ZZ/ZW sex determination mechanism (Becking *et al*., 2017). We sought to confirm female heterogamety by crossing females to sex-reversed males (which have female genotypes but male phenotypes), and assessing the sex ratio of the resulting progenies, which will differ depending on the sex chromosome system (XX neo-male × XX female yields all XX and therefore 100% female offspring; ZW neo-male × ZW female expected to produce 1/4 ZZ, 1/2 ZW, and 1/4 WW offspring, thus 75% female or 66.7% female depending on whether WW genotypes are viable). Surprisingly, we found that, at least in our focal population, sex is determined by an XX/XY system, suggesting a recent sex chromosome turnover. To test this hypothesis, we performed whole-genome sequencing. Consistent with a recent origin of an XX/XY sex determination system, we find evidence that the putative male-specific region is small relative to the whole genome, and we identified a male-specific, partial duplication of the androgenic gland hormone (AGH) gene, a rare example of a candidate sex-determining gene in a crustacean. In addition, although our study population does not appear to harbor current *Wolbachia* infections, we find genomic evidence of past infections. Overall, our results are consistent with the hypothesis that *Wolbachia* endosymbionts may have interfered with the evolution of sex determination in *T. rathkei*.

## Methods

### Animal collection and husbandry

We sampled wild isopods from Rice Creek Field Station (RCFS) at SUNY Oswego in Oswego, NY. We captured animals using a combination of methods. First, we haphazardly searched through leaf litter, logs, and rocks. We also used “potato traps”, made by carving out a 1-2 cm diameter core from a potato and placing it in the litter for 1-2 weeks. Finally, we constructed pitfall traps from plastic cups buried in the ground with the rim of the cup flush with the ground. The primary species captured were *Oniscus asellus* and *T. rathkei*, but we also captured *Philoscia muscorum*, *Hyloniscus riparius*, *Trichoniscus pusillus*, and occasionally *Cylisticus convexus*. Species identification was performed in the field and confirmed in the lab, where we also determined the phenotypic sex of specimens.

Isopods were housed in plastic food storage containers with holes in the lids for air exchange, on a substrate of moistened soil. Containers were checked twice weekly. Animals were fed carrots and dried leaves *ad libitum*. The photoperiod was kept on a schedule of 18:6 light hours: dark hours in the summer and 14:10 in the winter. We isolated ovigerous females in individual containers, and separated offspring from their mothers upon emergence from the marsupium.

We initially sexed offspring at six to eight weeks old, and separated males from females to prevent sibling mating. We then double-checked offspring sex at roughly two week intervals thereafter until four months of age to watch for individuals that might have shown late signs of sexual differentiation. Terrestrial isopods are known to store sperm from a single mating to fertilize future broods. Therefore, for experimental crosses we only used *T. rathkei* females that were born in the lab, separated from brothers as soon as they could be sexed, and which had not produced any offspring by 12 months of age.

### *Wolbachia* testing

We used PCR assays to test for *Wolbachia* presence in *T. rathkei* individuals. DNA was extracted from one or two legs, depending on the size of the animal. We ruptured the leg tissue μL deionized water along with a few 0.5 mm zirconia/silica beads (enough to cover the bottom of the tube) using a bead beater machine. Samples were lysed following a protocol of 2500 RPM for 10 seconds, followed by 4200 RPM for 10 seconds, and finally 4800 RPM for 10 seconds. The tube was then visually inspected to confirm the leg was sufficiently pulverized. We then transferred the lysate to a new tube, added 60 µL of a 5% Chelex® 100 molecular biology grade resin suspension, and incubated for 15 minutes at 100° C. After incubation, we centrifuged the extract at 16,000g for 3 minutes, and reserved 80 µL of supernatant for PCR testing. We confirmed successful DNA extraction using the mitochondrial primers HCO2198/LCO1490 (Folmer *et al*., 1994). We performed PCRs in 10 µL reactions, using 4.95 µL of molecular biology grade water, 2 µL NEB OneTaq Buffer, 1 µL of mixed dNTPS at a final concentration of 2mM for each dNTP, 1 µL of a 5 µM solution of each primer, and 0.05 µL of NEB OneTaq. For the mitochondrial primer set, PCR conditions included an initial denaturation of 94° C for 1 minute; 5 cycles of 94° C denaturation for 30s, 45° C annealing for 90s, and 68° C extension for 60s. The samples then underwent 35 cycles of 94° C for 30s, 51° C for 90s, and 68° C for 60s. This was followed by a final extension step of 68° C for 5 minutes. To test for *Wolbachia*, we performed PCR using *Wolbachia*-specific primers targeting the *wsp* (81f/691r) and *ftsZ* (ftsZf1/ftsZr1) genes (Werren John H. *et al*., 1995; Braig *et al*., 1998). We performed PCRs in 10 µL reactions, using 4.95 µL of molecular biology grade water, 2 µL NEB OneTaq Buffer, 1 µL of mixed dNTPs at a final concentration of 2mM for each dNTP, 1 µL of either *wsp* or *ftsZ* primers, and 0.05 µL of NEB OneTaq. PCR conditions contained an initial denaturation of 95°C for 5 minutes, followed by 36 cycles of 95° C for 60s, 54° C for 60s, and 68°C for 3 minutes. This was followed by a final extension step at 68° C for 10 minutes. Positive PCR tests would not necessarily be able to distinguish between a true infection and a copy of the *Wolbachia* genome horizontally integrated into the host genome, but the absence of a PCR product should be a reliable indicator that these *Wolbachia* sequences are not present (at least at detectable levels).

### Androgenic gland implantation and crosses

To test whether sex is determined by a ZZ/ZW or XX/XY system of sex determination in our population of *T. rathkei*, we performed crosses between females and experimentally sex-reversed neo-males. Juvenile female *T. rathkei* were implanted with live androgenic glands, according to (Becking *et al*., 2017). Male donors and female recipients were selected from large lab-reared broods with even (∼1:1) sex ratios. An adult male was sacrificed by decapitation, and live androgenic glands were dissected into Ringer solution (393 mM NaCl, 2 mM KCl, 2 mM CaCl_2_·2H_2_O, 2 mM NaHCO_3_). Female recipients were between 5 and 8 weeks old, an age at which males and females begin to become distinguishable by the appearance of external male genitalia, but at which sexual development is not complete. Occasionally, young males may still be mistaken for females at this stage, if the male genitalia are not sufficiently developed, and thus it is common for some fraction of the recipients in these experiments to carry male genotypes (Becking *et al*., 2017); nevertheless, it is important for transplant recipients to be as young as possible for sex reversal to be complete. The androgenic gland was injected using a pulled glass pipette into a hole pierced with a dissecting needle in the 6th or 7th segment of the juvenile female’s pereon. Recipients were isolated in a small plastic container with a moist paper towel for recovery and observation. Experimental animals were monitored for signs of male development. Any surviving animal that failed to develop male genitalia by 4 months post-implantation was considered to be a failed injection. After maturation, adult neo-males were placed in individual containers with 1-3 previously unmated females. Crosses were monitored twice weekly to check for signs of reproduction in females. Gravid females were then isolated into their own containers until parturition.

### Genome sequencing

We performed whole-genome sequencing using a combination of Illumina, PacBio, and Oxford Nanopore sequencing, with multiple sequencing samples of each sex (Supplementary Table 1). Because we expected the *T. rathkei* genome to be large, repetitive, and highly polymorphic, and because we expected to need to isolate DNA from multiple individuals, we established a partially inbred laboratory line using offspring from a single female collected from RCFS. We mated brothers and sisters from this female for two generations in the lab prior to collecting genetic samples from the third generation for sequencing. DNA was collected for sequencing using the Qiagen DNEasy Blood and Tissue Kit following the manufacturer’s instructions. DNA was quantified using the Qubit DNA Broad Range assay kit, and the A260/280 value was checked with a Nanodrop spectrophotometer. Samples were stored at −80°C prior to being shipped to the sequencing center. Illumina sequencing was performed at the State University of New York at Buffalo Genomics and Bioinformatics Core Facility.

For PacBio sequencing, we had to pool DNA from multiple individuals to obtain sufficient quantities of DNA for library preparation. We performed separate DNA extractions from three individuals of each sex as above. Then, we pooled the DNA from the three individuals of each sex and concentrated it using Ampure XP beads (Beckman-Coulter). Briefly, we washed the beads three times in molecular biology grade water, once in Qiagen buffer EB, and finally re-suspended the beads in their original buffer. We then added equal volumes of Ampure XP suspension to the DNA samples, mixed them on a shaker for 15 minutes, placed the tubes on a magnetic bead separator, and removed the supernatant. We washed the beads twice with 1.5 mL of 70% ethanol, and finally eluted the DNA samples in 30 µL of Qiagen buffer EB. Sequencing libraries were prepared and sequenced at the University of Delaware Sequencing & Genotyping Center on a PacBio RSII.

We also supplemented our PacBio dataset with Oxford Nanopore sequencing data. We isolated DNA from a single *T. rathkei* female and two separate males using a Qiagen DNEasy Kit as described above. We then performed sequencing on Oxford Nanopore Minion flowcells (R9.4) with the Rapid Sequencing Kit (SQK-RAD004) following the manufacturer’s instructions.

### Genome assembly

We performed a hybrid assembly combining the short- and long-read sequence data, closely mirroring the bioinformatics pipeline used to generate previously published isopod genome assemblies (Chebbi *et al*., 2019; Becking *et al*., 2019). We first removed adapters and trimmed the Illumina sequencing data using Trimmomatic v. 0.36 (Bolger *et al*., 2014); we removed leading and trailing bases, as well as internal windows of at least 4 bp, with a mean quality score of 5 or lower, and discarded any reads shorter than 36 bp after trimming. We then used SparseAssembler (Ye *et al*., 2012) to assemble the cleaned Illumina data from sample Mpool, This dataset was chosen because it had decent coverage (∼38x), was generated from a PCR-free sequencing library, and came from male samples (so that Y chromosome sequences would be present); additional male sequencing samples were excluded from this initial assembly to minimize the number of sequence polymorphisms that would be present in the data with additional samples. This first assembly was performed using two different kmer sizes (k=51 and k=61). After performing preliminary quality checks using Quast (Gurevich *et al*., 2013), we decided to proceed with the k=61 assembly, which had the longer total length and N50 (Supplementary Table 2). However, because we suspected the genome might still contain high levels of heterozygosity despite two generations of inbreeding, we used Redundans (Pryszcz and Gabaldón, 2016) to remove putative allelic contigs from the Illumina-only assembly; we set identity and overlap thresholds of 95%.

Prior to performing hybrid assembly, we used the short reads to correct sequencing errors in the long reads using FMLRC (Wang *et al*., 2018) with the default settings, except requiring a minimum count of 3 to consider a path (-m 3). PacBio and Oxford Nanopore reads derived from female isopods were corrected using Illumina sample Fpool, while long reads from male samples were corrected using sample Mpool.

We next performed hybrid assembly using DBG2OLC (Ye *et al*., 2016), which accepts a short-read assembly (rather than raw short-read sequence data) and long-read sequence data (in this case, our combined PacBio and Oxford Nanopore reads) as input. We tested out a range of different parameter values: from the Redundans-filtered assembly, we first removed contigs less than 100 bp or 200 bp; we tested kmer sizes of 17 and 19; for the kmer coverage threshold, we tried values of 2 and 5; and for the minimum overlap, we tried values of 10 and 30. We used an adaptive threshold of 0.01. These assemblies ranged in size from ∼5.2 Gb to 8.5 Gb; we selected three assemblies across the range of total sizes for further processing.

We next corrected errors in these assemblies, caused by the relatively high error rates in long-read sequence data. In the standard DBG2OLC pipeline, the resulting contigs are corrected using the contigs from the short-read assembly and from the long reads using Sparc (Ye and Ma, 2016); however, in our initial attempts, large portions of the assemblies went uncorrected, perhaps because we had relatively low-coverage long-read data. Therefore, instead we performed three rounds of error correction using Pilon (Walker *et al*., 2014), by mapping the trimmed Illumina sequence reads to each assembly using bbmap (first two rounds; Bushnell *et al*., 2017) and bwa mem (third round; with the parameters -A 1 -B 1 -O 1 -E 1 -k 11 -W 20 -d 0.5-L 6 for mapping to an error-prone assembly; Li, 2013).

Finally, we assessed the quality of each of the three candidate assemblies using BUSCO v.3.0.2 (Simão *et al*., 2015), with the arthropod reference gene set, and selected the assembly with the greatest number of BUSCO reference genes present for further analysis.

To remove contaminants from the final assembly, we generated blob plots using Blobtools v.1.0 (Laetsch and Blaxter, 2017). To accomplish this, we BLASTed all contigs against the NCBI nucleotide (nt) database using megablast (Morgulis *et al*., 2008) and against Uniprot reference proteomes using diamond blastx (Buchfink *et al*., 2015). We then removed any contigs that were identified as coming from plants, fungi, viruses, or bacteria, except for those matching *Wolbachia*.

### Genome annotation

We used RepeatModeler v.1.0.10, which uses RECON (Bao and Eddy, 2002), RepeatScout (Price *et al*., 2005), and Tandem Repeat Finder (Benson, 1999), to construct a custom repeat library for *T. rathkei*. Because we were unable to run RepeatModeler successfully using the full assembly, we ran RepeatModeler on a random subset 40% of the contigs; this should still successfully identify most repetitive elements in the genome as long as all repeat families are well represented in the subset. We then masked the assembly using RepeatMasker 4.0.7, and used the data generated by RepeatMasker on divergence between each individual repeat and the consensus sequence for that family to examine the history of transposable element activity in *T. rathkei* (Tarailo-Graovac and Chen, 2009).

We annotated coding sequences using the MAKER pipeline (Cantarel *et al*., 2008). We initially ran MAKER v2.31.9 using assembled transcript sequences (est2genome=1) from previously available data from one wild-caught male and one wild-caught female *T. rathkei* from the same population (Becking *et al*., 2017). This transcriptome was generated by assembling the male and female samples separately, each with two assemblers, Trinity v2.8.4 (Grabherr *et al*., 2011) and TransLiG v1.3 (Liu *et al*., 2019), and then merging and filtering the transcriptome assemblies with EvidentialGene v4 (Gilbert, 2019).This initial annotation also incorporated protein alignments against Uniprot-Swissprot (version March 2020),and the resulting output was used to train SNAP (Korf, 2004). To train AUGUSTUS (Stanke *et al*., 2006) we used the output from the BUSCO quality assessment described earlier. We then completed a final round of MAKER using the trained gene models, retaining the transcript and protein alignments from earlier as evidence.

### Development of sex-linked PCR markers

We used multiple approaches to develop male-specific, putatively Y-linked PCR markers. Initial attempts to perform a hybrid Illumina-PacBio genomic assembly with only male samples and then identify contigs with zero coverage in females were unsuccessful. We therefore developed a complementary approach by looking for male-specific k-mers using just the raw Illumina sequencing reads. We chose a value of k=21 because it should be large enough that most k-mers will not occur more than once in the genome sequence, yet small enough to minimize the impact of sequencing errors (Vurture *et al*., 2017). We used kmc v.3.1.0 (Kokot *et al*., 2017) to count all the canonical 21-mers in each of the Illumina sequencing datasets (in other words, each 21-mer and its reverse complement were considered to be the same k-mer during counting). We then searched for k-mers that occurred at least 8 times in the Mpool Illumina sequencing dataset and a total of at least 3 times combined across the lower coverage M2, M5, M6, and wildM samples, but which were completely absent from all female samples. We then extracted all Illumina sequence reads containing these candidate male-specific k-mers using mirabait v.4.0.2 (Chevreux *et al*., 1999), and assembled them using Spades v.3.11.1 (Bankevich *et al*., 2012). We also performed a reciprocal analysis looking for female-specific k-mers (which are not expected in an XX/XY system), searching for k-mers that occurred at least 8 times in the Fpool Illumina sequencing dataset, a total of at least 3 times combined across the F3, F4, and wildF samples, but which were completely absent from all male samples.

To test male-specificity of these contigs, we used PCR. We developed PCR primers for a subset of candidate male-specific contigs. To identify the best candidates, we first mapped raw sequencing reads from all male and female Illumina samples to the full genome sequence plus the candidate male-specific contigs, and identified contigs that had coverage in male samples but not female samples; we also avoided contigs that showed evidence of containing repeat elements, after BLAST searches against the whole genome assembly. We designed primers using PRIMER3 (Untergasser *et al*., 2012, p. 3). In these PCRs, primers were initially screened using template DNA from two male samples and two female samples; primers that showed evidence of sex specificity after this first PCR were re-tested using a larger number of samples. PCR primers were initially tested using a cycle of 98°C for 3 minutes, followed by 40 cycles of 98°C for 15s, 50°C for 35s, and 68°C for 60s; this was followed by a final extension step of 68°C for 10 minutes. For samples that did not amplify under this program, a gradient PCR was run to determine optimal annealing temperature. All PCRs were performed using the same recipe and reaction conditions as the *Wolbachia* PCRs described above.

We also identified open reading frames (ORFs) in these candidate male-specific contigs using Transdecoder v.4.0.0 (Haas and Papanicolaou, 2016), and annotated the ORFs using Trinotate v.3.1.1 (Bryant *et al*., 2017). Subsequently, we designed additional primers targeting one of the candidate ORFs (F: 5’-ATTCTTGACTCTCCCCACGA-3’; R: 5’-TCTCCAACTACGATTTCGTTAATT-3’).

## Results

### No *Wolbachia* and balanced sex ratios in *T. rathkei*

Among the 100+ individuals captured and tested between 2015 and 2017, no *T. rathkei* from RCFS conclusively tested positive for *Wolbachia*. This was not due to inadequate testing protocols; for instance, a captive population of *Porcellio laevis* housed in our lab shows nearly a 100% infection rate using the same methods (not shown). Approximately 150 *T. rathkei* broods were raised in the lab from either mated, wild-caught females or first-generation crosses. The mean and median brood sizes of this species in our lab were 27.1 and 22.5 offspring, respectively, and the vast majority of these broods had a balanced sex ratio (Supplementary Table 3). Thus, the prevalence of *Wolbachia* and other sex ratio distorters is at most very low in this population of *T. rathkei*. In addition, some wild-caught females produced broods even after several months to a year in isolation in the lab (Supplementary Table 3), confirming that this species is capable of long-term sperm storage.

### Crossing sex-reversed individuals indicates an XY sex determination system

Five juveniles implanted with androgenic glands survived to mature into males; they were crossed with virgin females from families with normal sex ratios. Each putative neo-male was paired with 2 to 3 females, and each female produced 1-3 broods of offspring. Two of these males sired broods with balanced sex ratios (not significantly different from the null hypothesis of a 1:1 ratio of males to females; Table 1). These males were likely individuals that would have developed into males even without the AG implantation, but were initially mis-identified as juvenile females probably due to incomplete sexual differentiation at that early stage. Thus, these crosses are uninformative with respect to the sex determination system. A similar rate of “failed” crosses was observed in a recent study following identical protocols in other isopod species, including species without any evidence of sex chromosome polymorphism, suggesting that sexing juveniles for AG implantation at these early stages is difficult due to the possibility of incomplete sexual differentiation (Becking *et al*., 2017). Three other males produced only female offspring, consistent with an XX/XY system (XX neo-male × XX female yields all XX and therefore 100% female offspring) but not a ZZ/ZW system (ZW neo-male × ZW female expected to produce 1/4 ZZ, 1/2 ZW, and 1/4 WW offspring, thus 75% female or 66.7% female depending on whether WW genotypes are viable; Table 1).

**Table 1.**
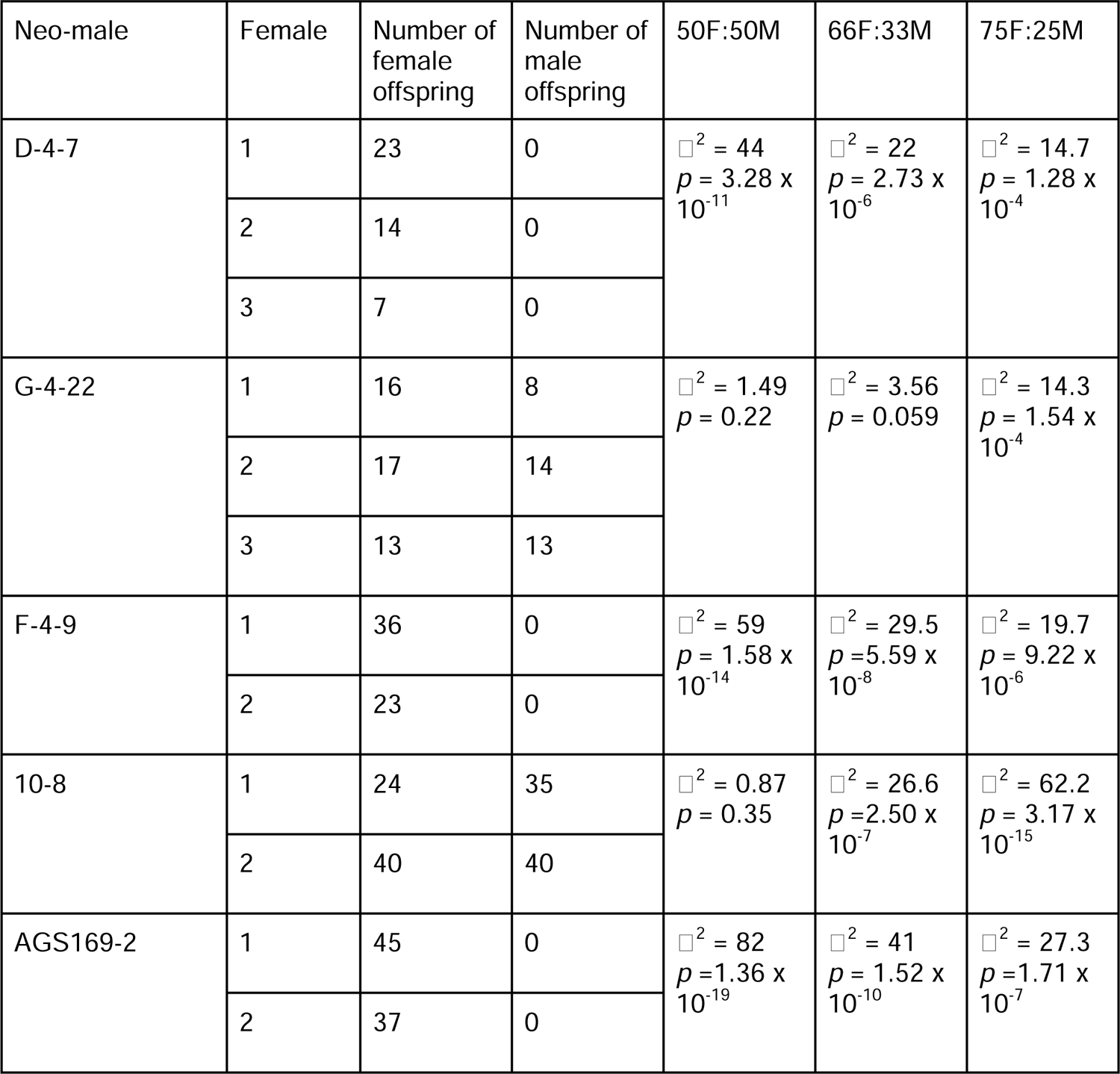
Sex ratios from crosses between putative neo-males (juvenile females implanted with an androgenic gland) and females. The last three columns give the results of chi-square tests testing whether the observed sex ratios (pooling the results for each male) are significantly different from predicted ratios of 1 female: 1 male, 2 female: 1 male, and 3 female: 1 male.

### Genome assembly

All sequencing data have been deposited in the NCBI SRA under the project accession number PRJNA633105 (sequencing runs are SRR11797353-SRR11797365, SRR4000573, and SRR4000567); the draft assembly has been deposited under accession number GCA_015478945.1. The draft genome assembly of *T. rathkei* is approximately 5.2 Gb in total length. The genome is highly repetitive, consisting of approximately 70% repetitive elements. Transposable elements constitute the largest repeat category, with LINEs, followed by DNA elements and LTRs, being the most represented (Figure 1). All repeat families seem to have a single divergence peak of around 7-10% (Figure 1).

**Figure 1.**
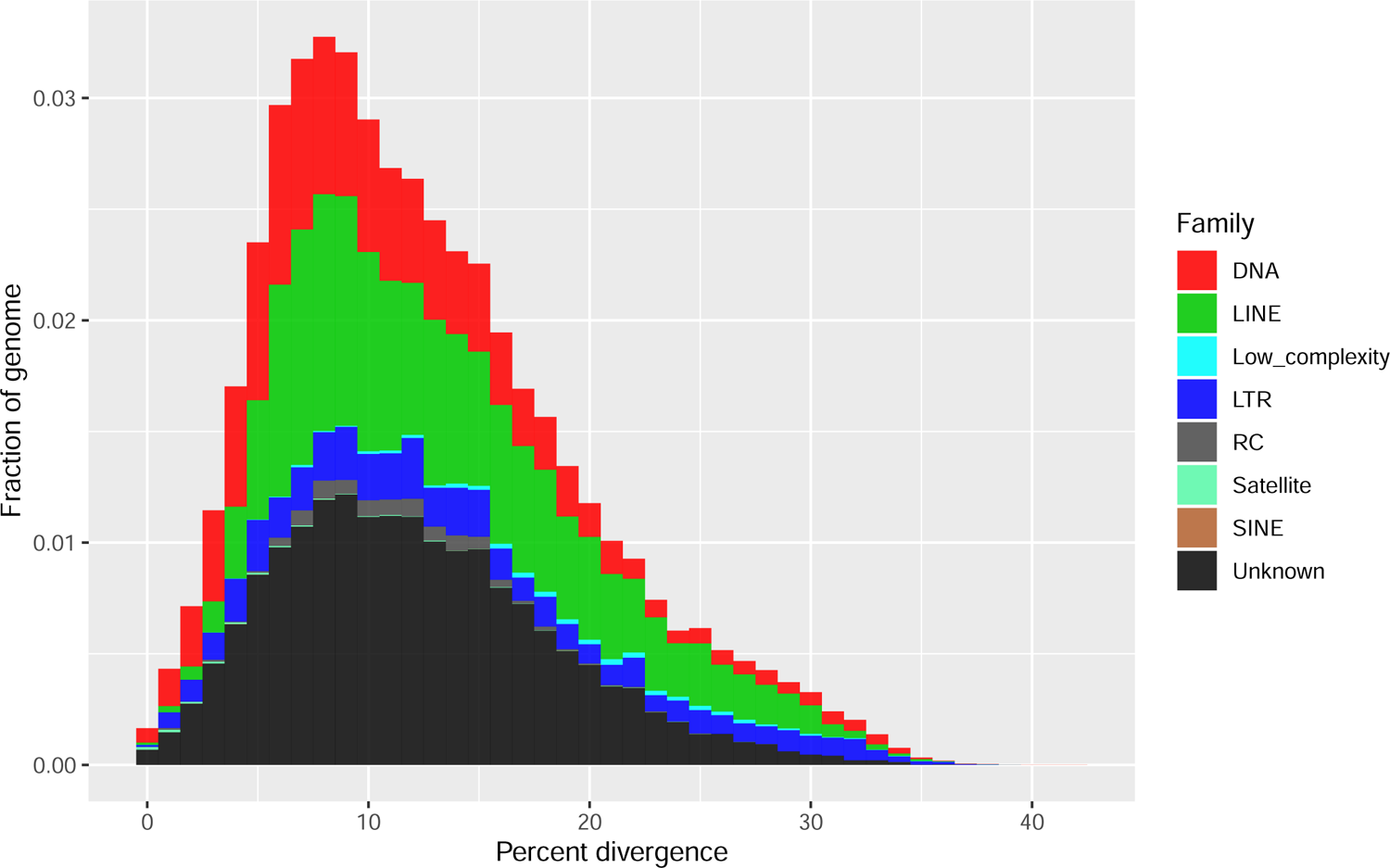
Distribution of divergence levels for repetitive elements in the *T. rathkei* genome.

Despite its large size, the draft assembly is likely only partially complete, with ∼25% of arthropod BUSCO genes missing (Table 2). For an independent assessment of assembly completeness, we also estimated the proportion of transcripts from the previously available transcriptome dataset that were present in the assembly. There were 15,805 transcripts assembled from that previously available transcriptome whose best hits in blastn searches against the NCBI nt database and diamond blastx searches against Uniprot-Swissprot were from other arthropods (and thus were unlikely to come from bacterial contaminants); of those, only 53% had nearly full-length matches in the genome (≥90% of the transcript length at ≥90% sequence identity), suggesting some missing data and/or remaining uncorrected sequencing errors in the draft assembly as well. At a more relaxed cutoff of at least 50% of the transcript having BLAST hits in the genome at ≥ 90% sequence identity, then 91% of transcripts appear to be represented somewhere in the assembly, suggesting that most transcripts are indeed present in the assembly but may be partial fragments rather than whole gene sequences.

**Table 2.** Assembly statistics for the *T. rathkei* draft genome.

|  |  |
| --- | --- |
| Total length | 5,181,251,014 bp |
| Number of contigs | 421,784 |
| N50 | 39,761 bp |
| GC content | 29.0% |
| Complete BUSCO genes | 533 single copy (51.9%); 39 duplicated (3.7%) |
| Fragmented BUSCO genes | 203 (19.0%) |
| Missing BUSCO genes | 271 (25.4%) |

We screened the *T. rathkei* genome for *Wolbachia* nuclear insertions by BLASTing the assembled contigs against a collection of *Wolbachia* genome sequences, and then BLASTing the matching regions against all representative bacterial genomes from RefSeq to rule out false positives. After this filtering step, we were left with 1,010 high confidence matches (best BLAST hit in a *Wolbachia* genome, e-value < 1 x 10^-6^) spread across 719 contigs, with a total length of ∼350 kb for the matching sequences (Supplementary Table 4), much smaller than a typical full *Wolbachia* genome of about 1 - 1.6 Mb on average (Sun *et al*., 2001). These may represent independent small insertions into the host genome, or one or more larger insertions that were subsequently broken up by mobile elements or other genomic rearrangements, or which assembled into separate contigs due to insufficient data. These likely horizontally acquired sequences were closely related to *Wolbachia* strain *wCon* from the isopod *Cylisticus convexus* (Badawi *et al*., 2018), the feminizing *Wolbachia* strain *wVulC*, and the *f* element of *A. vulgare* (Leclercq *et al*., 2016) (Figure 2).

**Figure 2.**
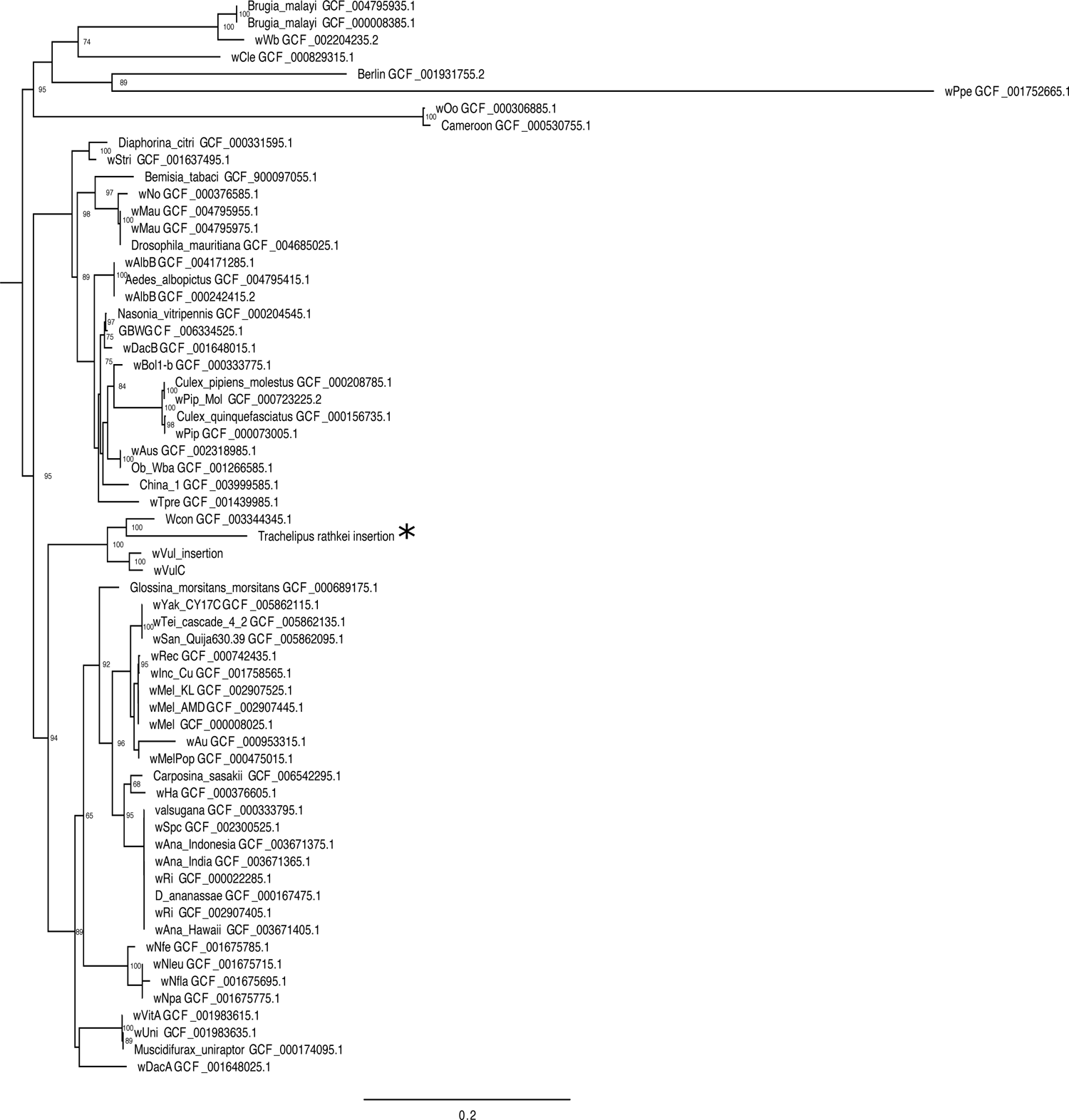
Phylogenetic tree showing the relationship of candidate horizontally transferred *Wolbachia* segments in the *T. rathkei* genome to other *Wolbachia* isolates. The *T. rathkei Wolbachia* insertion (indicated by the asterisk) is closely related to *Wolbachia* isolates from other isopods, and its closest relative is the wCon from *Cylisticus convexus.* Numbers by nodes indicate bootstrap support. Branch lengths represent average number of substitutions per site. The tree was generated by concatenating all candidate *Wolbachia* insertions in *T. rathkei* longer than 1,000 bp, along with the best-matching regions in the reference *Wolbachia* genomes (found with BLAST), aligning with MUSCLE v.3.8.31 (Edgar, 2004), filtering alignments with trimal v. 1.2rev59 (Capella-Gutiérrez *et al*., 2009), selecting a model using ModelTest-NG v. 0.1.6 (Darriba *et al*., 2020), and running the analysis in RAxML-NG v. 0.9.0 (Stamatakis, 2014) with 100 bootstrap replicates.

### Searching for candidate sex-determining genes

We identified ∼6.04 x 10^6^ 21-mers as potentially male-specific, suggesting there is a minimum of 6 Mb of male-specific sequence content in the genome. However, when we isolated the raw Illumina sequencing reads containing those 21-mers and assembled them, we obtained 89.4 Mb of assembled sequences, suggesting the male-specific region may be as large as ∼90 Mb, but still shares significant similarity with the X chromosome. Even if up to 90 Mb of sequence is partially sex-linked, this represents just 1.7% of the genome.

As a negative control, we also screened for k-mers specific to females. We identified 9.11 x 10^5^ female-specific 21-mers using the same criteria as our analysis for male-specific k-mers, or approximately 15% the number of male-specific k-mers. The observation that there are far more male-specific 21-mers than there are female-specific 21-mers in our dataset suggests that most of these are from real male-specific genomic sequences.

Of the initial 16 candidate Y-linked PCR markers designed from anonymous sequences, none showed the expected pattern of male-specific amplification in our early tests (Supplementary Table 5). This may be due to (1) the highly repetitive nature of the *T. rathkei* genome, despite our best efforts to target primers to non-repetitive sequences, or (2) low divergence between X- and Y-linked copies.

Because the candidate male-specific contigs were assembled from Illumina data only and thus short and fragmented, we were unable to screen them for annotated candidate sex-determining genes using the typical MAKER pipeline. However, we were able to identify open reading frames (ORFs) and annotate them like transcripts using Trinotate (Bryant *et al*., 2017). Three contigs in the male-specific assembly showed homology to the androgenic gland hormone (AGH) gene upon annotation, suggesting there may be a Y-linked duplication of the AGH gene. Therefore, we designed PCR primers specifically targeting one of the Y-linked AGH-like sequences (AGHY1 on NODE_44048_length_535, see Methods for primer sequences; same PCR cycling conditions as the other candidate sex-specific primers). These primers resulted in a PCR product of the expected size (195 bp) in all male samples screened (7/7), but not in any of the female samples (0/7), all of which were unrelated wild-caught individuals, confirming the male-specificity of this AGH allele.

This AGH sequence could be either a male-specific duplication of the AGH gene, or a Y-linked allele that has diverged from an X-linked copy (in other words, gametologs). To distinguish between these possibilities, we examined the sequencing depth of these genes and of other putatively single-copy genes (identified in the BUSCO analysis) in male and female Illumina sequencing data. If the male-specific AGH sequence is a gametolog of an X-linked sequence, we would expect the total sequencing depth of all AGH sequences (putative autosomal and putative Y-linked) to be the same in both the pooled male and pooled female samples, with the female sample having a higher average sequencing depth for the putative X-linked AGH sequences (since they would be homozygous for the X-linked gametologs, while males would be hemizygous for the X-linked gametologs). If, on the other hand, the male-specific AGH sequences are Y-linked duplicates, and not allelic to the other AGH sequences in our assembly, we would expect the shared autosomal AGH sequences to have similar sequencing depth in both male and female samples, and the combined sequencing depth of all AGH sequences (putative autosomal and putative Y-linked) would be higher in the male sample. Our results were consistent with the latter scenario, suggesting these are Y-linked duplicates rather than gametologs (Figure 3). Note that sequencing depth of AGHY1 and AGHY2, though much lower in the female sample than in the male sample, is still non-zero in the female sample, probably because of ambiguously mapped reads due to high similarity between the Y-linked and autosomal copies.

**Figure 3.**
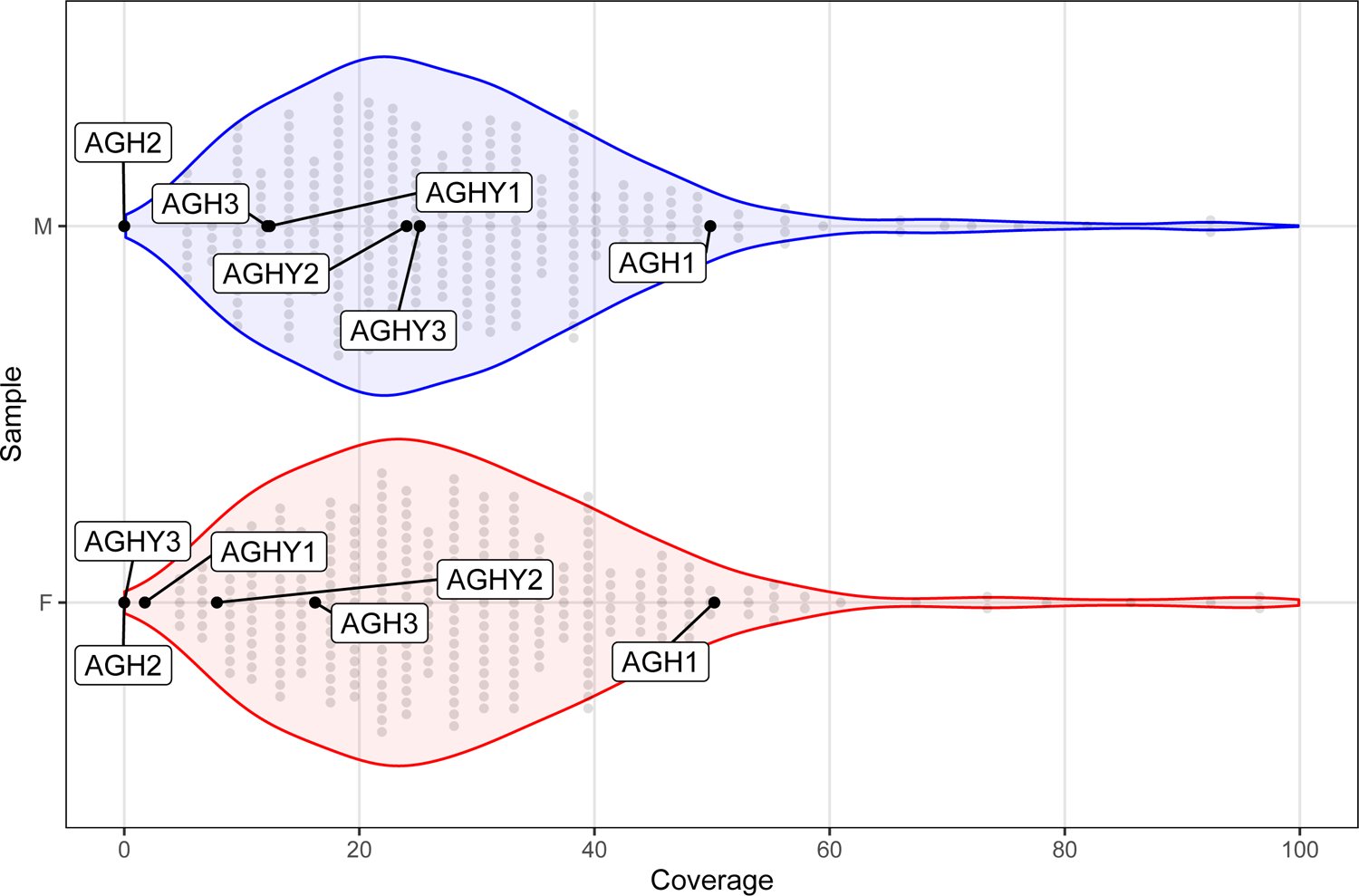
Distribution of sequencing depth for single-copy BUSCO genes in male and female Illumina sequencing datasets (M-pool and F-pool). Labeled dots indicate the sequencing depth for the different AGH copies in each sample.

Because the male-specific AGH sequences were found only in our Illumina data, we were unable to assemble them into long contigs, even after repeated attempts to assemble them individually with different assemblers and parameter values (not shown); all these contigs were ∼600 bp or less in length. Thus we are unable to determine whether these are complete duplicates of the whole gene, or fragments. Nevertheless, a phylogenetic analysis suggests that one of the Y-linked duplicates is a copy of the other, rather than an independent duplication of an autosomal copy, and based on branch lengths they are as divergent from one another as AGH orthologs in different species (Figure 5). In addition, these Y-linked copies seem to lack an intron that is present in the autosomal copies (Figure 4), suggesting they may have originated via retrotransposition. Note that AGHY1 and AGHY2 seem to be duplicates of the same region of the AGH gene (Figure 4), suggesting they are indeed two duplicates (the fact that both copies have appreciable sequencing depth in the pooled male sample, and are ∼20% divergent at the DNA sequence level, suggests that these are real duplicates and not assembly artifacts; moreover, they cannot be allelic sequences because all the males in the Illumina samples that they were assembled from were siblings, carrying the same Y chromosome). AGHY3, on the other hand, covers a different region of the AGH gene (Figure 4), so AGHY3 may not be an additional third duplicate, but may instead be a part of AGHY1 or AGHY2 that just assembled into a separate contig.

**Figure 4.**
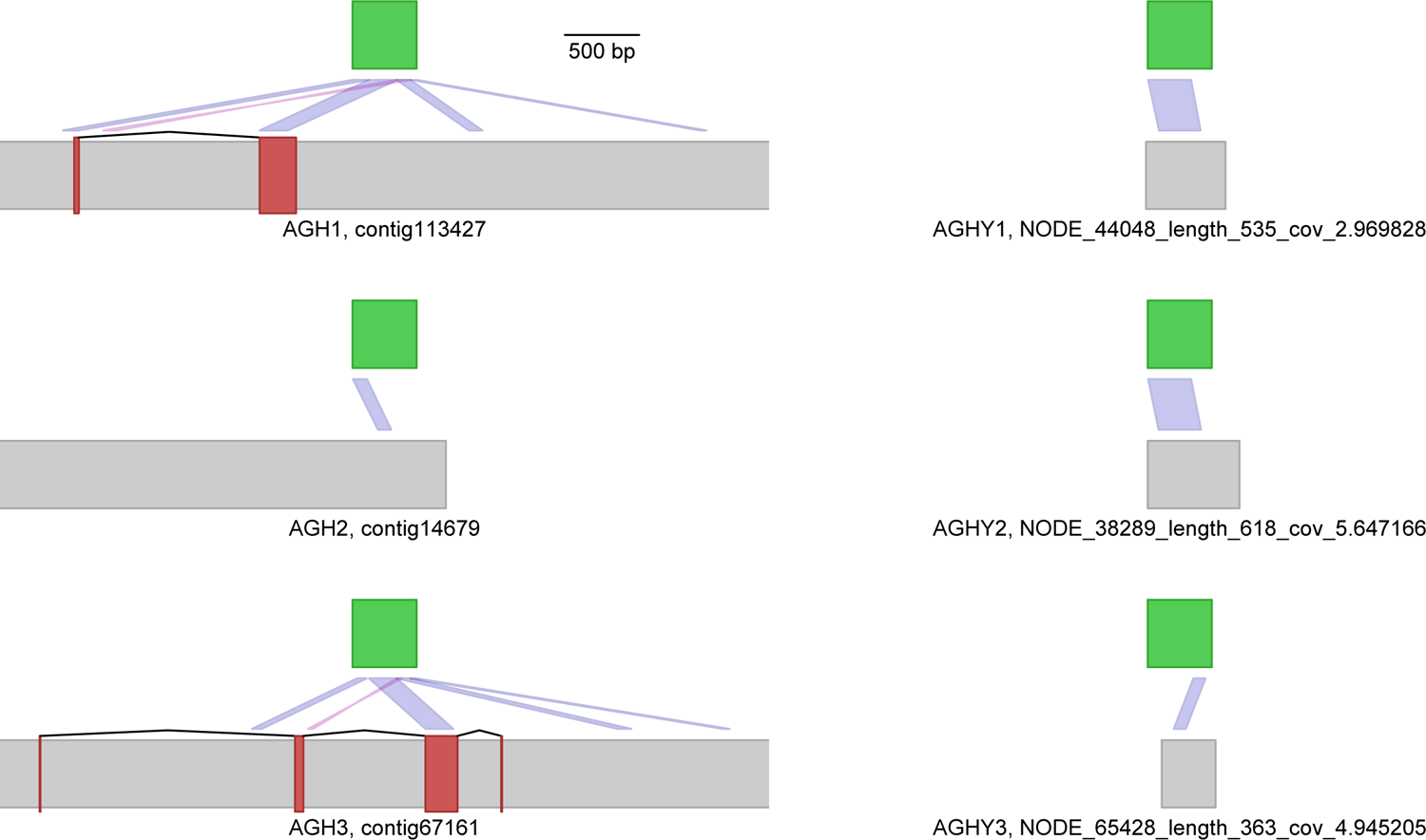
Possible duplicates of the androgenic gland hormone gene in the *T. rathkei* genome, including male-specific duplicates (on the right). The green bars represent the sequence of the expressed AGH sequence, assembled from previously available transcriptome data. Gray bars represent contigs in the draft genome assembly, and the pink bars on contigs represent exons annotated by MAKER. Dark blue segments connecting portions of the transcript to portions of contigs represent BLAST hits; light purple connector segments represent BLAST hits in reverse orientation. The incongruence between annotated exons and BLAST matches between the transcript and contigs suggests the annotation still contains some errors.

**Figure 5.**
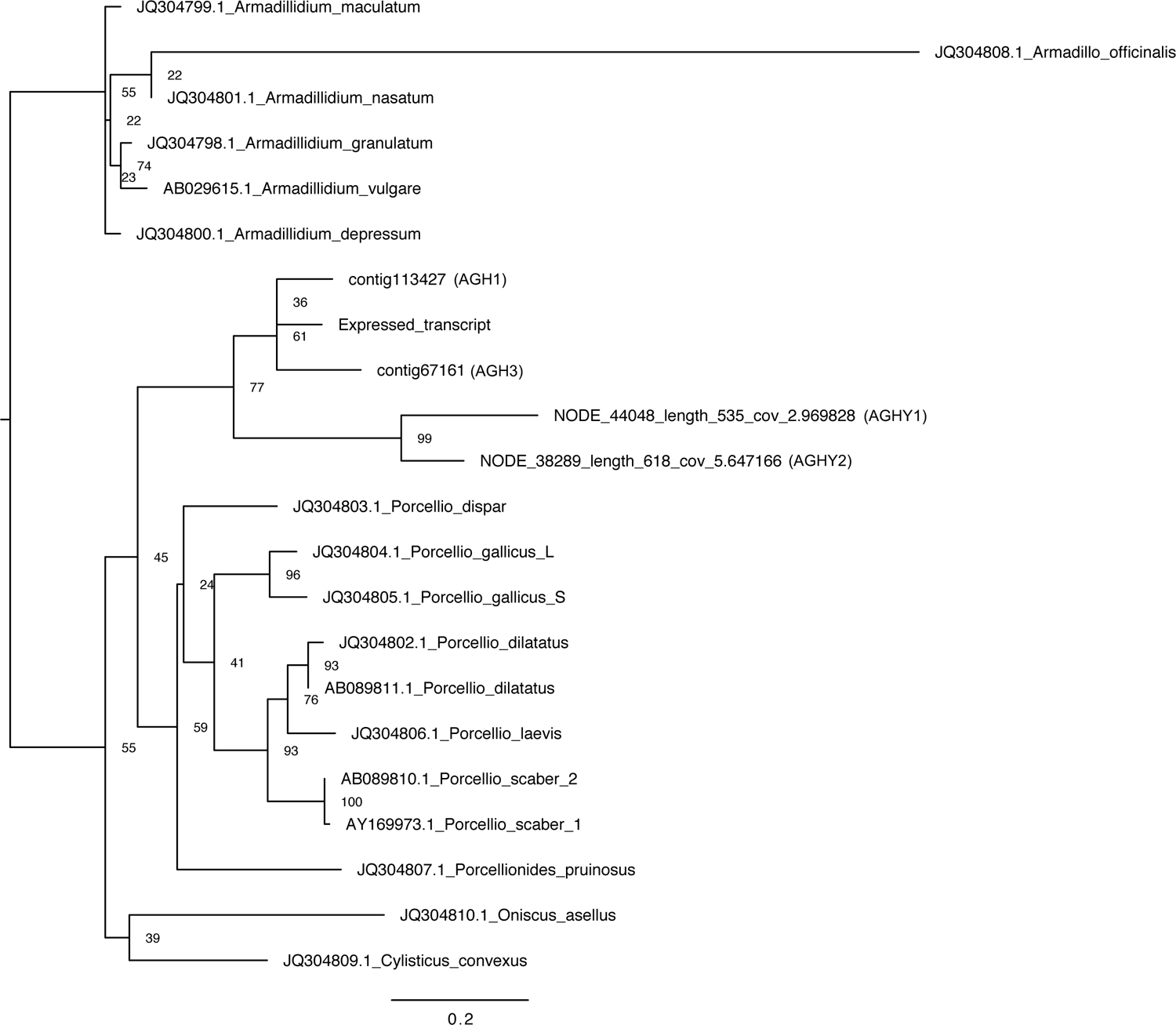
Phylogenetic tree showing relationships among AGH sequences from terrestrial isopods. AGH2 and AGHY3 are missing from this phylogeny because those sequences were omitted because of their short length. The tree was generated using all AGH-like sequences from *T. rathkei* of at least 100 bp, along with reference AGH nucleotide sequences from other species, aligning them with MUSCLE v. 3.8.31 (Edgar, 2004), selecting a model using ModelTest-NG v. 0.1.6 (Darriba *et al*., 2020), and running the analysis in RAxML-NG v. 0.9.0 (Stamatakis, 2014) with 100 bootstrap replicates.

We also find some evidence of additional autosomal duplicates of the androgenic gland hormone (AGH) gene. Two contigs in the full assembly contained annotated transcripts with at least partial homology to the expressed transcript identified as the AGH sequence, and a third contained no annotated genes but still showed high sequence similarity to AGH in BLAST searches. However, not all of the annotated exons in the first two copies matched the expressed transcript, and there were unannotated portions of the same contigs that did show sequence similarity to the transcript (Figure 4). Moreover, some of the matching portions of the assembled contigs had less than 90% sequence identity to the expressed transcript, and analysis of the sequencing depth of these regions reveals that one has very low coverage, suggesting it may be an assembly artifact (see below). Thus, we cannot rule out the possibility that some of these possible autosomal duplicates represent assembly and/or annotation artifacts. If they are real, these autosomal duplicates appear to be specific to *Trachelipus*, occurring after its divergence from *Porcellio* (Figure 5), but they may still be nonfunctional.

## Discussion

We have shown that, at least in our upstate New York population, sex determination in the terrestrial isopod *T. rathkei* is based on an XX/XY sex chromosome system. Two independent lines of evidence support this finding: first, crosses between females and sex-reversed neo-males yielded all female offspring (Table 1), consistent with an XX/XY system but not a ZZ/ZW system (Becking *et al*., 2017); second, we have identified PCR primers that only amplify a product in male samples, indicating the presence of a male-specific genomic region, i.e., a Y chromosome.

Our findings run counter to a previously published study showing evidence of female heterogamety in this species based on cytogenetics; in that study, female germ cells contained one set of unpaired chromosomes (presumably, the Z and W sex chromosomes), while male germ cells did not (Mittal and Pahwa, 1980). There are multiple possible explanations for this contradiction. First, it is possible that the previous study incorrectly identified the species of study specimens, as no information on identification is given in the paper; however, *T. rathkei* is relatively easy to distinguish from other cosmopolitan terrestrial isopod species by its five pairs of pleopodal “lungs” (most superficially similar species such as *Porcellio scaber* have only two pairs; Hatchett, 1947; Shultz, 2018). In addition, that study was published before feminizing *Wolbachia* was widely recognized in terrestrial isopods. It is therefore theoretically possible that the females used in that study carried an XY genotype but were feminized by *Wolbachia*, while the males in that study might have carried a YY genotype, perhaps resulting from a cross between an XY father and a sex-reversed XY or YY mother which failed to transmit *Wolbachia* (Becking *et al*., 2019).

Perhaps the most likely explanation is sex chromosome differences between populations. Indeed, this would not be unprecedented, as sex determination in terrestrial isopods is thought to evolve rapidly (Rigaud *et al*., 1997; Cordaux *et al*., 2011; Becking *et al*., 2017), and within-species sex chromosome polymorphisms are documented in a few other species. For instance, two subspecies of *Porcellio dilatatus*, *P. dilatatus dilatatus* and *P. dilatatus petiti* have XX/XY and ZZ/ZW systems, respectively (Juchault and Legrand, 1964; Legrand *et al*., 1974; Becking *et al*., 2017). In addition, multiple sex determining elements segregate in populations of the common pillbug *A. vulgare* (Juchault *et al*., 1992), including a novel W chromosome that resulted from the integration of an almost entire *Wolbachia* genome into the host genome (Leclercq *et al*., 2016). Outside terrestrial isopods, sex chromosome polymorphisms are also documented in a range of other arthropods and vertebrates (Orzack *et al*., 1980; Franco *et al*., 1982; Ogata *et al*., 2008; Rodrigues *et al*., 2013). *T. rathkei* is probably non-native in North America where this study was conducted (Jass and Klausmeier, 2000), as well as perhaps in India where the prior study on cytogenetics was done (Mittal and Pahwa, 1980). Given its cosmopolitan distribution, and the fact that other terrestrial isopods have moderate to high levels of genetic diversity (Romiguier *et al*., 2014), it might not be especially surprising for *T. rathkei* to harbor multiple polymorphic sex-determining loci. Nevertheless, the XX/XY system seems to be fixed, or at least the majority, in our population: multiple segregating sex-determining factors within a single populations usually result in sex-biased broods (Denholm *et al*., 1986; Basolo, 1994), including in other isopods like *A. vulgare* (Rigaud and Juchault, 1993); however, we observed only a few (out of 131) sex-biased broods from wild-caught gravid females or lab-reared females outside of our sex reversal experiments, and never observed any single-sex broods. Moreover, the male-specific primer pair we designed amplified successfully in all tested wild-caught males, and none of the wild-caught females, albeit with modest sample sizes. Hopefully future follow-up work can further characterize geographic variation in sex determination in this species.

Regardless of whether or not sex determination is polymorphic in our population of *T. rathkei*, we propose that the ZZ/ZW sex chromosomes in this species are more likely to be ancestral, and that the XX/XY system is derived. First, *T. rathkei* is nested within a clade that mostly consists of ZZ/ZW species (Becking *et al*., 2017). Moreover, the previous study finding a ZZ/ZW system in *T. rathkei* was based on the presence of heteromorphic sex chromosomes in female meiotic spreads (Mittal and Pahwa, 1980), suggesting that the Z and W have been diverging long enough to be cytogenetically distinguishable, in contrast to other isopods examined so far showing homomorphic sex chromosomes (Rigaud *et al*., 1997). In addition, the putative male-specific region of the *T. rathkei* genome does not contain any genes that are essential for male reproduction or spermatogenesis, since phenotypic males with an XX genotype still sired offspring in our sex-reversal experiments, and the male-specific region is only a small portion of the total genome, similar to other terrestrial isopods examined so far (Chebbi *et al*., 2019; Becking *et al*., 2019). This male-specific region is probably at least 6 Mb and has an upper size limit of around 90 Mb, but these estimates include sequences that retain high similarity to X-linked copies; indeed, most of the candidate male-specific primers we tested failed to show sex-specific amplification patterns. These observations suggest that this Y chromosome may be evolutionarily young, since it has not had time to accumulate major differences from the X. Given that we found genomic evidence of a past association with *Wolbachia* in this species and that infection by *Wolbachia* has been found in other *T. rathkei* populations (Cordaux *et al*., 2012), this scenario is consistent with the hypothesis that transitions in sex determination mechanisms may be triggered by *Wolbachia* and other endosymbionts that manipulate host reproduction (Rigaud *et al*., 1997; Cordaux *et al*., 2011). If other populations of *T. rathkei* with different sex determination mechanisms can be identified, it may be possible to leverage this system to further study the mechanisms and selective forces influencing transitions in sex determination mechanisms. In addition, studies of sex determination in a phylogenetic context involving other members of the family Trachelipodidae would shed further light on the origins of the X and Y chromosomes in *T. rathkei*.

The draft genome assembly of *T. rathkei* is especially large, at around 5.2 Gb, with approximately 29% GC content. The actual genome is likely to be even larger, given that ∼25% of the BUSCO arthropod orthologs were missing in our assembly. By comparison, genomes of pillbugs in the genus *Armadillidium* tend to be smaller at around 1.2 - 2 Gb in size (Chebbi *et al*., 2019; Becking *et al*., 2019), but other terrestrial isopods have genomes ranging to over 8 Gb (Gregory, 2020), and other crustacean relatives such as amphipods also have large genomes (Rees *et al*., 2007; Rivarola-Duarte *et al*., 2014; Kao *et al*., 2016), so *T. rathkei* is not out of the ordinary for this group.

The *T. rathkei* genome contains a large proportion of repetitive elements, in particular transposable elements (Figure 1). The most common transposable element families are LINEs, DNA elements, and LTRs, similar to *A. vulgare* and *A. nasatum* (Chebbi *et al*., 2019; Becking *et al*., 2019, 2020). The distribution of divergence values, with a single mode around 7-10% divergence (Figure 1), suggests that most repeat families expanded around the same time as previously shown in *A. vulgare* and *A. nasatum*; however, unlike in *A. vulgare*, *T. rathkei* shows no evidence of a second more recent burst in DNA element activity. Simple repeats also comprise a substantial portion of the genome; even manually looking through the assembled contigs reveals a high abundance of (TA)x repeats. It would be interesting to examine the repeat content of the male-specific portion of the genome. Unfortunately, however, we were only able to recover male-specific sequences from the short-read Illumina data, and this portion of the genome assembly is highly fragmented, precluding more detailed analysis. Hopefully, additional long-read sequencing data will allow us to examine transposable element dynamics in this area in the future.

We found many contigs with high similarity to the *Wolbachia* genome (Supplementary Table 4), even though we were unable to detect current *Wolbachia* infections in our population using PCR. This is not surprising given that horizontal transfers of Wolbachia DNA into host genomes is common (Dunning Hotopp, 2011), and that *Wolbachia* is relatively common in terrestrial isopods and arthropods in general (Cordaux *et al*., 2012; Pascar and Chandler, 2018; Medina *et al*., 2019) and has been found in other populations of *T. rathkei*. These *Wolbachia* insertions are closely related to other *Wolbachia* strains from isopods, including feminizing strains (Cordaux et al 2004, Leclercq et al 2016). This suggests that *T. rathkei* may have been infected with a feminizing *Wolbachia* strain in the past, even though no firm conclusion can be drawn solely from phylogenetic evidence. If so, it is conceivable that *Wolbachia* may have been involved in the sex chromosome turnover we characterized in *T. rathkei*, as previously hypothesized (Rigaud et al 1997, Cordaux and Gilbert 2017).

Male differentiation in terrestrial isopods is controlled by the androgenic gland hormone, AGH. AGH is a peptide hormone similar in structure to insulin, and is secreted by the androgenic gland (Martin *et al*., 1999). AGH expression is sufficient to transform juvenile female isopods into fertile males (Martin *et al*., 1999). Presumably, in wild-type males, the primary sex-determining signal triggers the differentiation of the androgenic glands during development, which then secretes AGH. Interestingly, the draft genome of *T. rathkei* contains multiple AGH-like sequences, unlike *A. vulgare*, which has a single copy (Chebbi et al 2019). While some of these may be assembly artifacts, there is evidence of at least two partial Y-linked sequences (assembled from Illumina sequencing reads containing male-specific k-mers), of which one was confirmed by PCR to be male-specific. These duplications seem to be specific to *T. rathkei* (Figure 5), though other members of the genus *Trachelipus* or the family Trachelipodidae have yet to be examined. Consistent with this, a past study found no evidence of any expressed AGH duplications in other terrestrial isopod species except *Porcellio gallicus* (Cerveau *et al*., 2014).

In many other species, novel sex chromosomes have arisen via duplication of a sex-determining gene. For instance, duplicates of the vertebrate gene *Dmrt1* have evolved into master sex-determining signals on the W and Y chromosomes, respectively, in the frog *Xenopus laevis* (Yoshimoto *et al*., 2008) and the medaka *Oryzias latipes* (Matsuda *et al*., 2002, 2007; Nanda *et al*., 2002), while a Y-linked duplicate of the anti-Müllerian hormone gene is a candidate master sex-determining gene in the teleost fish *Odontesthes hatcheri* (Hattori *et al*., 2012). The presence of Y-linked AGH copies in *T. rathkei*, and no other obvious open reading frames homologous to known sex determination or sex differentiation genes, makes these genes obvious candidates for the master male-determining signal in *T. rathkei*. Sex-specific genomic regions like the Y and W chromosomes are also expected to acquire sexually antagonistic alleles (van Doorn and Kirkpatrick, 2007, 2010; Charlesworth, 2017).Thus, if functional these duplicates might instead provide male fitness benefits rather than serving as a master male sex-determining gene. Unfortunately, we were unable to assemble full copies of these Y-linked AGH homologs because they only showed up in our Illumina data, not in our low-coverage long read data. Future deep sequencing using long reads should further clarify the molecular evolution of these genes. In addition, expression studies should determine which of these genes are expressed, in what tissues, and at what stages.

We have shown that the terrestrial isopod *T. rathkei* uses an XX/XY sex chromosome system, at least in upstate New York, in contrast to a past cytogenetic study suggesting a ZZ/ZW mechanism (Mittal and Pahwa, 1980). In line with this, whole-genome sequencing and follow-up PCRs demonstrate the existence of male-specific, Y-linked copies of the androgenic gland hormone gene in this species. These findings highlight the role of gene duplication in the evolution of sex chromosomes and they further establish terrestrial isopods as models to study the evolution of sex determination systems and the mechanisms underlying their transitions.

## Acknowledgments

We thank the editor and three anonymous reviewers for constructive comments on earlier drafts of this manuscript. We also appreciate computing time and assistance provided by the National Center for Genome Analysis Support at Indiana University, especially Tom Doak and Sheri Sanders. This research was funded by National Science Foundation grant NSF-DEB 1453298 to CHC. It was also supported in part by Lilly Endowment, Inc., through its support for the Indiana University Pervasive Technology Institute. This research is based upon work supported by the National Science Foundation under Grant Nos. DBI-1062432 2011, ABI-1458641 2015, and ABI-1759906 2018 to Indiana University. Any opinions, findings, and conclusions or recommendations expressed in this material are those of the authors and do not necessarily reflect the views of the National Science Foundation, the National Center for Genome Analysis Support, or Indiana University.

## Notes

https://github.com/chandlerlab/trachelipus_genome

### Competing Interest Statement

The authors have declared no competing interest.

