## Supplementary Tables for "Evolutionary transition to XY sex chromosomes associated with Y-linked duplication of a male hormone gene in a terrestrial isopod"

**Supplementary Table 1.** *T. rathkei* samples used for genomic sequencing.

| Sample name | Sex | Notes | Platform/ read length | Total data (Gb) | Ref. | Accession Number |
| --- | --- | --- | --- | --- | --- | --- |
| M-pool | M | Pool of three brothers, lab-reared | Illumina, 2x100 | 191.6 | This study | SRR11797365 |
| F-pool | F | Pool of three sisters, lab-reared | Illumina, 2x100 | 191.9 | This study | SRR11797364 |
| M2 | M | One male, lab-reared | Illumina, 2x250 | 22.3 | This study | SRR11797360 |
| M5 | M | One male, lab-reared | Illumina, 2x250 | 20.6 | This study | SRR11797359 |
| M6 | M | One male, lab-reared | Illumina, 2x250 | 23.6 | This study | SRR11797358 |
| F3 | F | One female, lab-reared | Illumina, 2x250 | 131.5 | This study | SRR11797357 |
| F4 | F | One female, lab-reared | Illumina, 2x250 | 26.8 | This study | SRR11797356 |
| Wild-M | M | One male, wild-caught | Illumina, 2x100 | 14.1 | (Chandler *et al.*, 2015) | SRR4000567 |
| Wild-F | F | One female, wild-caught | Illumina, 2x100 | 14.4 | (Chandler *et al.*, 2015) | SRR4000573 |
| PB-F1 | F | Pool of three sisters, lab-reared | PacBio | 22.2 | This study; (Peccoud *et al.*, 2017) | SRR11797355 |
| PB-F2 | F | Pool of three sisters, lab-reared | PacBio | 0.2 | This study; (Peccoud *et al.*, 2017) | SRR11797354 |
| PB-M | M | Pool of three brothers, lab-reared | PacBio | 3.9 | This study; (Peccoud *et al.*, 2017) | SRR11797353 |
| ONT-F | F | One female | ONT | 0.4 | This study | SRR11797363 |
| ONT-M1 | M | One male | ONT | 3.9 | This study | SRR11797362 |
| ONT-M2 | M | One male | ONT | 3.3 | This study | SRR11797361 |

**Supplementary Table 2.** Assembly statistics for initial genome assemblies performed with only short-read data. k=61 was selected as the optimal short-read assembly for further hybrid assembly with long-read data using DBG2OLC.

|  | k = 51 | k = 61 |
| --- | --- | --- |
| # contigs (>= 0 bp) | 62,285,342 | 53,534,813 |
| # contigs (>= 500 bp) | 750,475 | 966,258 |
| # contigs (>= 1000 bp) | 148,561 | 227,660 |
| # contigs (>= 5000 bp) | 850 | 1088 |
| Total length (>= 0 bp) | 5,689,084,525 | 5,960,525,287 |
| Total length (>= 500 bp) | 626,371,545 | 846,993,489 |
| Total length (>= 1000 bp) | 224,744,531 | 347,268,680 |
| Total length (>= 5000 bp) | 7,240,744 | 7,187,453 |
| Largest contig (bp) | 73,394 | 28,080 |
| GC (%) | 33.6 | 33.6 |
| N50 (bp) | 817 | 877 |

**Supplementary Table 3.** Sex ratios of *T. rathkei* broods reared in the lab. Wild-caught refers to a mated female captured in the wild and brought back to the lab to give birth. Only broods with at least 10 offspring are shown.

| Cross type | Mother ID | Father ID | Brood number | Year born | Male | Female | Total Offspring | % Female | *ᵪ*^2^ | *p* |
| --- | --- | --- | --- | --- | --- | --- | --- | --- | --- | --- |
| Wild-caught | 8 | unknown | 1 | 2016 | 14 | 12 | 26 | 46.2 | 0.15 | 0.695 |
| Wild-caught | 10 | unknown | 1 | 2016 | 19 | 15 | 34 | 44.1 | 0.47 | 0.493 |
| Wild-caught | 11 | unknown | 1 | 2016 | 11 | 11 | 22 | 50.0 | 0 | 1 |
| Wild-caught | 13 | unknown | 1 | 2016 | 18 | 12 | 30 | 40.0 | 1.2 | 0.273 |
| Wild-caught | 14 | unknown | 1 | 2016 | 26 | 15 | 41 | 36.6 | 2.95 | 0.086 |
| Wild-caught | 14 | unknown | 2 | 2016 | 7 | 10 | 17 | 58.8 | 0.53 | 0.467 |
| Wild-caught | 17 | unknown | 1 | 2016 | 23 | 28 | 51 | 54.9 | 0.49 | 0.484 |
| Wild-caught | 17 | unknown | 2 | 2016 | 8 | 10 | 18 | 55.6 | 0.22 | 0.637 |
| Wild-caught | 18 | unknown | 1 | 2016 | 6 | 12 | 18 | 66.7 | 2 | 0.157 |
| Wild-caught | 19 | unknown | 1 | 2016 | 15 | 13 | 28 | 46.4 | 0.14 | 0.705 |
| Wild-caught | 20 | unknown | 1 | 2016 | 7 | 6 | 13 | 46.2 | 0.08 | 0.782 |
| Wild-caught | 21 | unknown | 1 | 2016 | 20 | 18 | 38 | 47.4 | 0.11 | 0.746 |
| Wild-caught | 21 | unknown | 2 | 2016 | 4 | 8 | 12 | 66.7 | 1.33 | 0.248 |
| Wild-caught | 22 | unknown | 1 | 2016 | 17 | 14 | 31 | 45.2 | 0.29 | 0.59 |
| Wild-caught | 25 | unknown | 1 | 2016 | 19 | 12 | 31 | 38.7 | 1.58 | 0.209 |
| Wild-caught | 26 | unknown | 1 | 2016 | 9 | 42 | 51 | 82.4 | 21.35 | 3.82e-06** |
| Wild-caught | 26 | unknown | 2 | 2016 | 2 | 13 | 15 | 86.7 | 8.07 | 4.51e-03** |
| Wild-caught | 28 | unknown | 1 | 2016 | 12 | 16 | 28 | 57.1 | 0.57 | 0.45 |
| Wild-caught | 28 | unknown | 2 | 2016 | 8 | 5 | 13 | 38.5 | 0.69 | 0.405 |
| Wild-caught | 29 | unknown | 1 | 2016 | 12 | 20 | 32 | 62.5 | 2 | 0.157 |
| Wild-caught | 29 | unknown | 2 | 2016 | 8 | 9 | 17 | 52.9 | 0.06 | 0.808 |
| Wild-caught | 30 | unknown | 1 | 2016 | 20 | 26 | 46 | 56.5 | 0.78 | 0.376 |
| Wild-caught | 32 | unknown | 1 | 2016 | 5 | 16 | 21 | 76.2 | 5.76 | 0.016* |
| Wild-caught | 34 | unknown | 1 | 2016 | 12 | 11 | 23 | 47.8 | 0.04 | 0.835 |
| Wild-caught | 35 | unknown | 1 | 2016 | 5 | 5 | 10 | 50.0 | 0 | 1 |
| Wild-caught | 39 | unknown | 1 | 2016 | 6 | 5 | 11 | 45.5 | 0.09 | 0.763 |
| Wild-caught | 42 | unknown | 1 | 2016 | 8 | 10 | 18 | 55.6 | 0.22 | 0.637 |
| Wild-caught | 45 | unknown | 1 | 2016 | 3 | 7 | 10 | 70.0 | 1.6 | 0.206 |
| Wild-caught | 46 | unknown | 1 | 2016 | 17 | 24 | 41 | 58.5 | 1.2 | 0.274 |
| Wild-caught | 47 | unknown | 1 | 2016 | 15 | 8 | 23 | 34.8 | 2.13 | 0.144 |
| Wild-caught | 47 | unknown | 2 | 2017 | 33 | 30 | 63 | 47.6 | 0.14 | 0.705 |
| Wild-caught | 47 | unknown | 3 | 2017 | 27 | 20 | 47 | 42.6 | 1.04 | 0.307 |
| Wild-caught | 48 | unknown | 2 | 2017 | 23 | 25 | 48 | 52.1 | 0.08 | 0.773 |
| Wild-caught | 77 | unknown | 1 | 2016 | 6 | 8 | 14 | 57.1 | 0.29 | 0.593 |
| Wild-caught | 83 | unknown | 1 | 2016 | 6 | 4 | 10 | 40.0 | 0.4 | 0.527 |
| Wild-caught | 83 | unknown | 2 | 2017 | 15 | 20 | 35 | 57.1 | 0.71 | 0.398 |
| Wild-caught | 89 | unknown | 1 | 2016 | 12 | 7 | 19 | 36.8 | 1.32 | 0.251 |
| Wild-caught | 106 | unknown | 1 | 2016 | 8 | 10 | 18 | 55.6 | 0.22 | 0.637 |
| Wild-caught | 107 | unknown | 1 | 2016 | 5 | 5 | 10 | 50.0 | 0 | 1 |
| Wild-caught | 115 | unknown | 1 | 2016 | 8 | 4 | 12 | 33.3 | 1.33 | 0.248 |
| Wild-caught | 116 | unknown | 1 | 2016 | 7 | 9 | 16 | 56.3 | 0.25 | 0.617 |
| Wild-caught | 119 | unknown | 1 | 2017 | 24 | 26 | 50 | 52.0 | 0.08 | 0.777 |
| Wild-caught | 121 | unknown | 1 | 2016 | 11 | 8 | 19 | 42.1 | 0.47 | 0.491 |
| Wild-caught | 122 | unknown | 1 | 2016 | 6 | 12 | 18 | 66.7 | 2 | 0.157 |
| Wild-caught | 123 | unknown | 1 | 2016 | 7 | 5 | 12 | 41.7 | 0.33 | 0.564 |
| Wild-caught | 128 | unknown | 1 | 2016 | 3 | 8 | 11 | 72.7 | 2.27 | 0.132 |
| Wild-caught | 128 | unknown | 2 | 2017 | 24 | 17 | 41 | 41.5 | 1.2 | 0.274 |
| Wild-caught | 130 | unknown | 1 | 2016 | 11 | 2 | 13 | 15.4 | 6.23 | 0.013* |
| Wild-caught | 132 | unknown | 1 | 2016 | 9 | 12 | 21 | 57.1 | 0.43 | 0.513 |
| Wild-caught | 136 | unknown | 1 | 2016 | 5 | 6 | 11 | 54.5 | 0.09 | 0.763 |
| Wild-caught | 151 | unknown | 1 | 2017 | 23 | 25 | 48 | 52.1 | 0.08 | 0.773 |
| Wild-caught | 161 | unknown | 1 | 2017 | 15 | 15 | 30 | 50.0 | 0 | 1 |
| Wild-caught | 161 | unknown | 2 | 2017 | 17 | 24 | 41 | 58.5 | 1.2 | 0.274 |
| Wild-caught | 163 | unknown | 1 | 2016 | 23 | 14 | 37 | 37.8 | 2.19 | 0.139 |
| Wild-caught | 163 | unknown | 2 | 2017 | 28 | 47 | 75 | 62.7 | 4.81 | 0.028* |
| Wild-caught | 163 | unknown | 3 | 2017 | 4 | 18 | 22 | 81.8 | 8.91 | 2.84e-03** |
| Wild-caught | 165 | unknown | 1 | 2017 | 21 | 24 | 45 | 53.3 | 0.2 | 0.655 |
| Wild-caught | 165 | unknown | 2 | 2017 | 28 | 16 | 44 | 36.4 | 3.27 | 0.07 |
| Wild-caught | 167 | unknown | 1 | 2016 | 22 | 21 | 43 | 48.8 | 0.02 | 0.879 |
| Wild-caught | 167 | unknown | 2 | 2017 | 7 | 5 | 12 | 41.7 | 0.33 | 0.564 |
| Wild-caught | 168 | unknown | 1 | 2017 | 4 | 9 | 13 | 69.2 | 1.92 | 0.166 |
| Wild-caught | 169 | unknown | 1 | 2016 | 24 | 20 | 44 | 45.5 | 0.36 | 0.546 |
| Wild-caught | 169 | unknown | 2, 3 | 2016 | 30 | 40 | 70 | 57.1 | 1.43 | 0.232 |
| Wild-caught | 171 | unknown | 1 | 2017 | 22 | 21 | 43 | 48.8 | 0.02 | 0.879 |
| Wild-caught | 171 | unknown | 2 | 2017 | 19 | 21 | 40 | 52.5 | 0.1 | 0.752 |
| Wild-caught | 171 | unknown | 3 | 2018 | 22 | 23 | 45 | 51.1 | 0.02 | 0.881 |
| Wild-caught | 174 | unknown | 1 | 2016 | 21 | 20 | 41 | 48.8 | 0.02 | 0.876 |
| Wild-caught | 174 | unknown | 2 | 2017 | 17 | 26 | 43 | 60.5 | 1.88 | 0.17 |
| Wild-caught | 175 | unknown | 1 | 2016 | 25 | 17 | 42 | 40.5 | 1.52 | 0.217 |
| Wild-caught | 175 | unknown | 2 | 2016 | 18 | 17 | 35 | 48.6 | 0.03 | 0.866 |
| Wild-caught | 175 | unknown | 3 | 2017 | 28 | 29 | 57 | 50.9 | 0.02 | 0.895 |
| Wild-caught | 177 | unknown | 1 | 2016 | 20 | 11 | 31 | 35.5 | 2.61 | 0.106 |
| Wild-caught | 177 | unknown | 2 | 2016 | 15 | 18 | 33 | 54.5 | 0.27 | 0.602 |
| Wild-caught | 178 | unknown | 1 | 2017 | 21 | 19 | 40 | 47.5 | 0.1 | 0.752 |
| Wild-caught | 178 | unknown | 2 | 2017 | 10 | 17 | 27 | 63.0 | 1.81 | 0.178 |
| Wild-caught | 178 | unknown | 3, 4 | 2018 | 19 | 22 | 41 | 53.7 | 0.22 | 0.639 |
| Wild-caught | 179 | unknown | 1 | 2017 | 19 | 18 | 37 | 48.6 | 0.03 | 0.869 |
| Wild-caught | 179 | unknown | 2 | 2017 | 15 | 21 | 36 | 58.3 | 1 | 0.317 |
| Wild-caught | 179 | unknown | 3 | 2018 | 10 | 10 | 20 | 50.0 | 0 | 1 |
| Wild-caught | 180 | unknown | 1, 2 | 2017 | 18 | 14 | 32 | 43.8 | 0.5 | 0.48 |
| Wild-caught | 182 | unknown | 1 | 2017 | 7 | 10 | 17 | 58.8 | 0.53 | 0.467 |
| Wild-caught | 182 | unknown | 2 | 2017 | 7 | 7 | 14 | 50.0 | 0 | 1 |
| Wild-caught | 182 | unknown | 3 | 2018 | 20 | 15 | 35 | 42.9 | 0.71 | 0.398 |
| Wild-caught | 184 | unknown | 1, 2 | 2017 | 27 | 36 | 63 | 57.1 | 1.29 | 0.257 |
| Wild-caught | 185 | unknown | 2 | 2017 | 8 | 10 | 18 | 55.6 | 0.22 | 0.637 |
| Wild-caught | 186 | unknown | 2 | 2017 | 8 | 10 | 18 | 55.6 | 0.22 | 0.637 |
| Wild-caught | 187 | unknown | 1, 2 | 2017 | 11 | 13 | 24 | 54.2 | 0.17 | 0.683 |
| Wild-caught | 191 | unknown | 1, 2 | 2017 | 39 | 33 | 72 | 45.8 | 0.5 | 0.48 |
| Wild-caught | 192 | unknown | 1 | 2017 | 19 | 21 | 40 | 52.5 | 0.1 | 0.752 |
| Wild-caught | 192 | unknown | 2 | 2017 | 6 | 11 | 17 | 64.7 | 1.47 | 0.225 |
| Wild-caught | 192 | unknown | 3 | 2018 | 10 | 6 | 16 | 37.5 | 1 | 0.317 |
| Wild-caught | 193 | unknown | 1 | 2017 | 8 | 2 | 10 | 20.0 | 3.6 | 0.058 |
| Wild-caught | 194 | unknown | 2 | 2017 | 4 | 15 | 19 | 78.9 | 6.37 | 0.012* |
| Wild-caught | 196 | unknown | 2 | 2017 | 17 | 21 | 38 | 55.3 | 0.42 | 0.516 |
| Wild-caught | 196 | unknown | 3 | 2018 | 14 | 15 | 29 | 51.7 | 0.03 | 0.853 |
| Wild-caught | 197 | unknown | 1 | 2017 | 19 | 25 | 44 | 56.8 | 0.82 | 0.366 |
| Wild-caught | 197 | unknown | 2 | 2017 | 9 | 11 | 20 | 55.0 | 0.2 | 0.655 |
| Wild-caught | 199 | unknown | 1 | 2017 | 9 | 7 | 16 | 43.8 | 0.25 | 0.617 |
| Wild-caught | 199 | unknown | 2 | 2017 | 7 | 6 | 13 | 46.2 | 0.08 | 0.782 |
| Wild-caught | 201 | unknown | 1, 2 | 2017 | 11 | 12 | 23 | 52.2 | 0.04 | 0.835 |
| Wild-caught | 201 | unknown | 3 | 2018 | 7 | 7 | 14 | 50.0 | 0 | 1 |
| Wild-caught | 202 | unknown | 1 | 2017 | 7 | 13 | 20 | 65.0 | 1.8 | 0.18 |
| Wild-caught | 202 | unknown | 2 | 2018 | 5 | 9 | 14 | 64.3 | 1.14 | 0.285 |
| Wild-caught | 203 | unknown | 1 | 2017 | 7 | 14 | 21 | 66.7 | 2.33 | 0.127 |
| Wild-caught | 204 | unknown | 1 | 2017 | 17 | 13 | 30 | 43.3 | 0.53 | 0.465 |
| Wild-caught | 204 | unknown | 2 | 2017 | 15 | 14 | 29 | 48.3 | 0.03 | 0.853 |
| Wild-caught | 205 | unknown | 1 | 2017 | 21 | 13 | 34 | 38.2 | 1.88 | 0.17 |
| Wild-caught | 205 | unknown | 2 | 2017 | 6 | 11 | 17 | 64.7 | 1.47 | 0.225 |
| Wild-caught | 207 | unknown | 1, 2 | 2017 | 37 | 32 | 69 | 46.4 | 0.36 | 0.547 |
| Wild-caught | 208 | unknown | 1 | 2017 | 8 | 9 | 17 | 52.9 | 0.06 | 0.808 |
| Wild-caught | 208 | unknown | 2 | 2017 | 10 | 14 | 24 | 58.3 | 0.67 | 0.414 |
| Wild-caught | 211 | unknown | 1, 2 | 2017 | 27 | 13 | 40 | 32.5 | 4.9 | 0.027* |
| Lab cross | 10-F1 | 18-M1 | 1 | 2019 | 24 | 24 | 48 | 50.0 | 0 | 1 |
| Lab cross | 13-F1 | 107-M1 | 1 | 2018 | 8 | 9 | 17 | 52.9 | 0.06 | 0.808 |
| Lab cross | 151-M1 | 178-F1 | 1 | 2019 | 21 | 35 | 56 | 62.5 | 3.5 | 0.061 |
| Lab cross | 163-F1 | 163-M1 | 1 | 2019 | 34 | 42 | 76 | 55.3 | 0.84 | 0.359 |
| Lab cross | 169-F1 | 179-M1 | 1 | 2019 | 16 | 18 | 34 | 52.9 | 0.12 | 0.732 |
| Lab cross | 171-F1 | 162-M1 | 1 | 2018 | 5 | 8 | 13 | 61.5 | 0.69 | 0.405 |
| Lab cross | 177-F1 | 177/162-M1 | 1 | 2019 | 5 | 13 | 18 | 72.2 | 3.56 | 0.059 |
| Lab cross | 177-F2 | 177/162-M2 | 1 | 2019 | 10 | 20 | 30 | 66.7 | 3.33 | 0.068 |
| Lab cross | 199-F1 | 199-M1 | 1 | 2018 | 7 | 9 | 16 | 56.3 | 0.25 | 0.617 |
| Lab cross | 199-F1 | 199-M1 | 2 | 2018 | 6 | 12 | 18 | 66.7 | 2 | 0.157 |
| Lab cross | 199-F1 | 199-M1 | 3 | 2019 | 3 | 11 | 14 | 78.6 | 4.57 | 0.033* |
| Lab cross | 205-F1 | 205-M1 | 1 | 2018 | 19 | 10 | 29 | 34.5 | 2.79 | 0.095 |
| Lab cross | 208-F1 | 208-M1 | 1 | 2018 | 20 | 24 | 44 | 54.5 | 0.36 | 0.546 |
| Lab cross | 211-F1 | 211-M1 | 1 | 2019 | 15 | 4 | 19 | 21.1 | 6.37 | 0.012* |
| Lab cross | 34-F1 | 174-M1 | 1 | 2019 | 17 | 20 | 37 | 54.1 | 0.24 | 0.622 |
| Lab cross | 47-F1 | 47-M1 | 1 | 2018 | 38 | 37 | 75 | 49.3 | 0.01 | 0.908 |
| Lab cross | 47-F1 | 47-M1 | 2 | 2019 | 19 | 14 | 33 | 42.4 | 0.76 | 0.384 |
| Lab cross | 48-M1 | 48-F1 | 1 | 2019 | 26 | 27 | 53 | 50.9 | 0.02 | 0.891 |
| Lab cross | 83-F1 | 203-M1 | 1 | 2019 | 5 | 5 | 10 | 50.0 | 0 | 1 |
| Lab cross | 9-M1 | 132-F1 | 1 | 2018 | 25 | 10 | 35 | 28.6 | 6.43 | 0.011* |

**Supplementary Table 4.** Top 100 regions of the *Trachelipus rathkei* assembly matching one or more *Wolbachia* genomes.

| Contig | Matching *Wolbachia* genome | E-value | Bitscore | Length | Percent Identity |
| --- | --- | --- | --- | --- | --- |
| contig41205:46995-50826 | NZ_QPIP01000040.1 | 0 | 5802 | 3898 | 92.894 |
| contig21655:23176-27650 | NZ_CP016430.1 | 0 | 4834 | 4092 | 85.973 |
| contig82518:11250-14109 | NZ_QPIP01000225.1 | 0 | 4420 | 2894 | 93.988 |
| contig24014:19607-22010 | NZ_QPIP01000012.1 | 0 | 3807 | 2393 | 95.153 |
| contig82518:31954-35917 | NZ_CAOH01000056.1 | 0 | 3714 | 4061 | 80.325 |
| contig22648:52901-55829 | NZ_QPIP01000225.1 | 0 | 3610 | 2955 | 87.174 |
| contig60599:36848-52880 | NZ_CP041215.1 | 0 | 3606 | 4456 | 77.536 |
| contig111166:29090-31313 | NZ_MNCG01000080.1 | 0 | 2438 | 2274 | 83.641 |
| contig101838:5953-13532 | NZ_CP041215.1 | 0 | 2355 | 2341 | 81.888 |
| contig104492:6893-11374 | NZ_CP041215.1 | 0 | 1894 | 2067 | 79.971 |
| contig24014:16199-17706 | NZ_MUJL01000059.1 | 0 | 1819 | 1384 | 88.584 |
| contig140396:3608-5757 | NZ_QPIP01000011.1 | 0 | 1807 | 1695 | 83.481 |
| contig6767:28422-30540 | NZ_QPIP01000011.1 | 0 | 1807 | 1695 | 83.481 |
| contig62661:89289-93752 | NZ_CP041215.1 | 0 | 1794 | 1975 | 79.747 |
| contig54256:25557-27714 | NZ_QPIP01000011.1 | 0 | 1784 | 1705 | 82.991 |
| contig10767:48224-50433 | NZ_NSDS01000079.1 | 0 | 1773 | 2019 | 79.693 |
| contig83347:6124-8304 | NZ_CP016430.1 | 0 | 1707 | 2000 | 79.1 |
| contig8667:11430-15085 | NZ_CP015510.2 | 0 | 1693 | 2461 | 75.457 |
| contig82518:22137-23924 | NZ_QPIP01000225.1 | 0 | 1681 | 1160 | 92.5 |
| contig4956:73158-74195 | NZ_CAOH01000048.1 | 0 | 1635 | 1039 | 94.706 |
| contig54255:38034-40169 | NZ_QPIP01000011.1 | 0 | 1634 | 1708 | 81.03 |
| contig28080:13103-16290 | NZ_CP015510.2 | 0 | 1625 | 2450 | 74.735 |
| contig54256:19275-20850 | NZ_LRUH01000003.1 | 0 | 1618 | 1434 | 85.146 |
| contig41156:47828-49356 | NZ_QPIP01000070.1 | 0 | 1611 | 1570 | 83.312 |
| contig120126:10876-13379 | NZ_KB223538.1 | 0 | 1608 | 1371 | 86.287 |
| contig6767:22276-23850 | NZ_LRUH01000003.1 | 0 | 1604 | 1441 | 84.733 |
| contig4956:78999-82475 | NZ_VCEG01000122.1 | 0 | 1592 | 1355 | 85.83 |
| contig4865:65822-67934 | NZ_QPIP01000120.1 | 0 | 1591 | 1221 | 88.37 |
| contig4956:77436-78667 | NZ_QPIP01000116.1 | 0 | 1537 | 1206 | 88.557 |
| contig110376:12951-16136 | NZ_CP015510.2 | 0 | 1481 | 2536 | 73.226 |
| contig45554:30664-32437 | NZ_QPIP01000040.1 | 0 | 1466 | 970 | 93.505 |
| contig26619:11904-13635 | NZ_KB223538.1 | 0 | 1457 | 1494 | 81.124 |
| contig12055:11602-13784 | NZ_QPIP01000093.1 | 0 | 1446 | 1213 | 86.562 |
| contig55499:34817-37014 | NZ_KB223538.1 | 0 | 1446 | 1236 | 86.084 |
| contig120015:7288-11014 | NZ_CP015510.2 | 0 | 1439 | 2357 | 73.568 |
| contig56420:32935-36636 | NZ_CP015510.2 | 0 | 1405 | 2481 | 72.229 |
| contig57015:7061-9509 | NZ_CAOH01000065.1 | 0 | 1397 | 1300 | 83.692 |
| contig97794:44314-47449 | NZ_CP015510.2 | 0 | 1366 | 2461 | 72.694 |
| contig43090:31693-34077 | NZ_CP015510.2 | 0 | 1361 | 1863 | 76.436 |
| contig59781:1759-2780 | NZ_CAOH01000048.1 | 0 | 1343 | 1043 | 90.604 |
| contig57015:21936-26306 | NZ_CAOH01000065.1 | 0 | 1337 | 1259 | 83.4 |
| contig36754:35171-36510 | NZ_KB223538.1 | 0 | 1330 | 1402 | 80.243 |
| contig581:4071-5440 | NZ_NSDS01000070.1 | 0 | 1269 | 1152 | 84.549 |
| contig42215:18348-19410 | NZ_NSDS01000070.1 | 0 | 1257 | 1080 | 85.741 |
| contig131151:4311-5431 | NZ_NSDS01000070.1 | 0 | 1247 | 1139 | 84.197 |
| contig6767:8936-12307 | NZ_NSDS01000070.1 | 0 | 1242 | 1154 | 83.882 |
| contig47840:6928-9012 | NZ_NSDS01000070.1 | 0 | 1236 | 1145 | 83.843 |
| contig108259:12794-14948 | NZ_NSDS01000070.1 | 0 | 1228 | 1138 | 83.743 |
| contig110377:9706-11307 | NZ_RWIK01000003.1 | 0 | 1224 | 1581 | 77.293 |
| contig97486:21655-25569 | NZ_CP041215.1 | 0 | 1200 | 1590 | 76.541 |
| contig66402:15885-18640 | NZ_LK055284.1 | 0 | 1199 | 1815 | 74.49 |
| contig10737:33945-36112 | NZ_NSDS01000070.1 | 0 | 1194 | 1146 | 83.333 |
| contig107168:4876-8422 | NZ_LK055284.1 | 0 | 1191 | 1822 | 74.424 |
| contig56531:7008-8374 | NZ_LSYY01000126.1 | 0 | 1179 | 810 | 91.728 |
| contig25813:9937-11304 | NZ_LSYY01000126.1 | 0 | 1177 | 811 | 91.615 |
| contig8667:23939-25563 | NZ_RWIK01000003.1 | 0 | 1177 | 1651 | 75.954 |
| contig56531:9367-10734 | NZ_LSYY01000126.1 | 0 | 1172 | 811 | 91.492 |
| contig17820:18438-21324 | NZ_CTEH01000015.1 | 0 | 1169 | 829 | 91.435 |
| contig79792:23503-24741 | NZ_LSYY01000126.1 | 0 | 1166 | 812 | 91.379 |
| contig79792:26192-27559 | NZ_LSYY01000126.1 | 0 | 1163 | 811 | 91.245 |
| contig28080:22749-24395 | NZ_RWIK01000003.1 | 0 | 1162 | 1561 | 77.13 |
| contig98868:31083-33755 | NZ_CTEH01000006.1 | 0 | 1154 | 2240 | 71.607 |
| contig56531:4623-6003 | NZ_LSYY01000126.1 | 0 | 1150 | 824 | 90.413 |
| contig72521:1-909 | NZ_CP041215.1 | 0 | 1095 | 932 | 85.73 |
| contig109514:3774-5787 | NZ_NSDS01000070.1 | 0 | 1070 | 996 | 83.735 |
| contig136622:7675-8872 | NZ_CAOH01000065.1 | 0 | 1049 | 963 | 84.112 |
| contig32215:18066-20042 | NZ_CTEH01000006.1 | 0 | 1040 | 2005 | 71.521 |
| contig98868:20772-22347 | NZ_RWIK01000003.1 | 0 | 1028 | 1637 | 74.221 |
| contig2827:52033-53374 | NZ_CTEH01000015.1 | 0 | 1027 | 1351 | 76.166 |
| contig42215:16743-18227 | NC_021084.1 | 0 | 1022 | 946 | 83.932 |
| contig58604:31642-34836 | NZ_CTEH01000006.1 | 0 | 1022 | 2450 | 69.918 |
| contig84414:19934-21197 | NZ_CP041215.1 | 0 | 1020 | 922 | 84.382 |
| contig97794:50638-52378 | NZ_RWIK01000003.1 | 0 | 1017 | 1380 | 76.304 |
| contig131151:5506-7007 | NZ_KB223533.1 | 0 | 1003 | 935 | 83.316 |
| contig103716:10622-12081 | NZ_NSDS01000070.1 | 0 | 998 | 1026 | 81.189 |
| contig136941:1093-1804 | NZ_QPIP01000070.1 | 0 | 980 | 714 | 91.036 |
| contig4956:87666-89184 | NZ_CAOH01000037.1 | 0 | 952 | 750 | 88 |
| contig113691:13388-16226 | NZ_CP041215.1 | 0 | 940 | 1375 | 75.345 |
| contig62357:26051-29274 | NC_010981.1 | 0 | 930 | 2607 | 68.7 |
| contig22648:60781-63093 | NZ_CP041215.1 | 0 | 915 | 1236 | 75.971 |
| contig90834:10305-13093 | NZ_CTEH01000006.1 | 0 | 900 | 1594 | 72.271 |
| contig5091:23161-24553 | NZ_QPIP01000017.1 | 0 | 898 | 1403 | 74.626 |
| contig24485:37434-40025 | NZ_CP041215.1 | 0 | 889 | 1709 | 72.147 |
| contig98868:22871-23971 | NZ_RWIK01000003.1 | 0 | 884 | 992 | 80.444 |
| contig28080:19443-20340 | NZ_RWIK01000003.1 | 0 | 864 | 882 | 81.859 |
| contig20306:1-723 | NZ_CP011148.1 | 0 | 850 | 729 | 85.597 |
| NODE_4409_length_2226_cov_13.672390:1-980 | NZ_CAOH01000065.1 | 0 | 847 | 745 | 85.235 |
| contig8667:22312-23414 | NZ_RWIK01000003.1 | 0 | 837 | 972 | 79.835 |
| contig64403:13606-14996 | NZ_KB223533.1 | 0 | 830 | 667 | 87.706 |
| contig51480:5191-10342 | NZ_CP041215.1 | 0 | 825 | 1196 | 75.084 |
| contig110377:12182-13291 | NZ_RWIK01000003.1 | 0 | 820 | 984 | 78.455 |
| contig67097:19963-22152 | NZ_CP041215.1 | 0 | 815 | 1132 | 75.883 |
| NODE_1677_length_3365_cov_23.488464:190-1484 | NZ_CAOH01000066.1 | 0 | 792 | 963 | 78.712 |
| contig42215:13837-14533 | NZ_RWIK01000007.1 | 0 | 775 | 668 | 85.629 |
| contig21865:12389-13705 | NZ_KB223533.1 | 0 | 769 | 679 | 85.272 |
| contig55499:23126-24227 | NZ_CP034335.1 | 0 | 750 | 735 | 82.313 |
| contig4956:83945-85499 | NZ_CAOH01000037.1 | 0 | 736 | 727 | 82.256 |
| contig122193:35489-36065 | NZ_NSDS01000070.1 | 0 | 725 | 578 | 87.889 |
| contig82518:38729-39602 | NZ_LSYY01000140.1 | 0 | 720 | 885 | 78.418 |

**Supplementary Table 5.** Initial set of candidate primers for Y chromosome markers. None of the primer pairs from this initial set produced the expected patterns of male-specific amplification.

| ID | Forward primer | Reverse primer | Expected product size |
| --- | --- | --- | --- |
| TrYJa19_3 | GGGACACTGTTGTGAGAGCA | CGCCAACAACCAGATGCG | 160 |
| TrYJa19_21 | CTCTCACGTTCGGAACCACT | TCACTTTGGTTTTGCATGTCGA | 118 |
| TrYJa19_9 | TTCCTTTCCACCCCTATTTGCA | GCAAAGTGCTTACATGATTGCC | 100 |
| TrYJa19_19 | CGTGTCATTTTCTGTCAGCGA | CCCAAGGCAATGAAGCTGA | 114 |
| TrYJa19_4 | ACATCAGCAATCGCATCGT | TCGAGGTTGCGGATTCTGT | 120 |
| TrYJa19_5 | AGGCCCTATTAGGAAGACCCA | AAGGCCTGCTACCATGCG | 104 |
| TrYJa19_22 | AGCCTTGTGTGACCTTTCGA | CCTCACAGGCGGGACAAA | 145 |
| TrYJa19_16 | TGGTGTCATTTTTAACGGCAGA | GCAGGCAGTTTGTTTGCGA | 140 |
| TrYJa19_18 | GAGCAACAAGAATTCCTCGTTG | TCCGAGAACAAGTAAATCGTGC | 100 |
| TrYJa19_20 | CGCAGCCGCTGAAGGTAT | CACTTCCCATCAAAGTGTCCA | 176 |
| TrYJa19_1 | TGCAACTCATGGATTGTAAGCA | GGACACTGTCCCGTTCAAAAC | 101 |
| TrYJa19_7 | TTTGGCGTGGTTATGGCA | GTGTCCCAGTTATCAGGAACCT | 108 |
| TrYJa19_10 | TGGTGGCATTCTTTTCTCTGC | ACCACTAAGGGTTACCGCT | 237 |
| TrYJa19_6 | CCCAGGATTCTTCGAGCCC | TGCAGAGTGATCCCCGGT | 234 |
| TrYJa19_17 | GAAGGAGTCAGGCGCGTT | GCTTCTTGCGTTAGACGAAAT | 236 |
| TrYJa19_8 | CATCTTCTGGATGCGGCT | TGGCATCGTGTGTTCCGA | 228 |
